# Balancing memory fidelity and representational stability in the female mouse accessory olfactory bulb

**DOI:** 10.1101/2021.10.27.466073

**Authors:** Michal Yoles-Frenkel, Stephen D. Shea, Ian G. Davison, Yoram Ben-Shaul

## Abstract

Sensory systems must balance the value of efficient coding schemes against the need to update specific memorized representations without perturbing other memories. Here we describe a unique solution to this challenge that is implemented by the vomeronasal system (VNS) to encode and remember multiple conspecific individuals as part of the Bruce Effect (BE). In the BE, exposure of a pregnant female mouse to the odors of an unfamiliar male leads to failure of the pregnancy (‘pregnancy block’) via the VNS. Following mating and sensory exposure, however, the female becomes protected from a pregnancy block by the stud individual. While this form of natural learning has been proposed to depend on changes in the representation of his odors in her accessory olfactory bulb (AOB), a key VNS structure, there are no direct comparisons of *in vivo* sensory responses before and after imprinting. It has further been suggested that these changes simply render the AOB insensitive to stud odors. However, the combinatorial odor code used by the AOB and the significant overlap in the odor composition of different males means that silencing responses to one individual is likely to degrade responses to others, posing potential problems for more general sensory encoding. To identify the neuronal correlates of learning in the context of the BE, we recorded extracellular responses of AOB neurons *in vivo* in mated and unmated female mice upon controlled presentation of urinary chemosignals, including urine from both the stud and males of a distinct strain. We find that while initial sensory responses in the AOB (within a timescale required to guide social interactions) remain stable, responses to extended stimulation (as required for eliciting the pregnancy block) display selective attenuation of stud-responsive neurons. Based on our results, we propose a model that reconciles the formation of strong, selective memories with the need to sustain robust representational bandwidth by noting a distinction between the representations of brief and extended stimuli. This temporal disassociation allows attenuation of slow-acting endocrine processes in a stimulus-specific manner, without compromising consistent ongoing representations of stimuli that guide behavior.

## Introduction

A key function of learning is to associate stimuli with appropriate behavioral responses. Often, learning involves incremental changes integrated over multiple exposures to a stimulus, but sometimes, it can be much more rapid. A dramatic example of the latter is behavioral imprinting. This unique form of learning usually coincides with a crucial life event which stimulates acquisition of a memory during a single exposure [1]. Typically, imprinting involves linking a stereotyped and robust response to a complex stimulus that is *a priori* unknown. Among the best-known and most widely-replicated examples of imprinting in mammals is the Bruce effect (BE), first described and repeatedly shown in various forms in mice [2-4] and later demonstrated or implicated in other species as well [5-10].

In mice, the BE involves two related components. The first is the innate sensory-mediated pregnancy *block*, in which exposure of a pregnant female mouse to an unfamiliar male mouse, or to his urine [11], during the first 72 hours post mating [2], leads to pregnancy failure. The second component, and the focus of this work, involves learning. Specifically, during mating the female forms a memory of the stud male, lasting several weeks, so that his odors will no longer elicit pregnancy failure. In the context of the BE, imprinting involves an interaction between individual recognition, and the hypothalamic circuits that mediate basic physiological and endocrine responses. A series of studies has shown that both pregnancy block and the learning that selectively circumvents it involve sensory detection by the vomeronasal system (VNS), a chemosensory system devoted to processing cues from other organisms [12-14]. More specifically, it was suggested that activation of the VNS by unfamiliar male chemosignals would trigger a cascade of events involving the amygdala and hypothalamic dopamine [15, 16] neurons that block secretion of prolactin, which in turn is required for maintenance of pregnancy during the initial stages [17]. Importantly, and consistent with a key role of the VNS in the BE, several studies have demonstrated the capacity of the VNS to represent information about conspecific individuals [18-21].

What changes occur during mating that prevent the stud male from inducing a pregnancy block of his own litter? The prevailing theory is that mating leads to selective silencing of stud responsive neurons in the first brain relay of the VNS, the accessory olfactory bulb (AOB) [3, 22-24]. According to this idea, known as the ‘negative template hypothesis’ (NTH), odors associated with the stud male will no longer be activated powerfully enough to trigger the hormonal cascade that leads to pregnancy block. While very elegant, this solution faces two fundamental difficulties. First, complete silencing of activity in response to the stud would render the female anosmic to her mate, at least with respect to vomeronasal mediated recognition, a scenario which appears maladaptive. Second, cross-talk between representations of different stimuli is likely to cause interference between the learned odors and other biologically relevant cues. AOB representations are combinatorial [21, 25-31] so that single AOB neurons often respond to a range of stimuli. As a consequence, AOB neuronal ensembles associated with distinct stimuli, sometimes with highly divergent ethological implications, may display extensive overlap. This implies that silencing, or mere attenuation of stud activated neurons will alter representations of multiple of stimuli in a complex manner. These considerations address a broader issue, namely the balance between plasticity and stability in neuronal representations. Clearly, plasticity is an essential feature of sensory processing, but when it alters sensory representations at an early processing stage such as the AOB, and compromises the stability of representations of multiple stimuli, it can complicate and even confound the readout of neuronal activity. How then can plasticity in the context of the BE take place without sacrificing representational stability?

One potential solution to this puzzle is suggested by a recent *in vitro* study investigating the effects of mating on AOB neuronal activity [32]. The study observed a novel form of plasticity in which mating-activated projection neurons (mitral tufted cells - MTCs) show reduced excitability. Critically, the reduced excitability was not apparent during a single current injection stimulus, but rather appeared only following an extended stimulation sequence. This raised the possibility that mating-induced plasticity is not expressed as a simple and direct silencing of activity, but rather as an impaired ability to sustain robust activation for extended periods. This observation suggests a scenario in which mating would change the stud’s ability to affect the slow endocrine processes required for induction of pregnancy failure [3], without altering the initial responses to sensory stimuli.

While *in vitro* data are suggestive, direct measurements of how sensory learning affects AOB representations in the intact brain are lacking. The key goal of the present work was to test, for the first time, the effects of mating on sensory representations at the level of the AOB *in vivo*. Consistent with the prior *in vitro* work [32], we found that prolonged exposure to the stud male’s urine evoked responses that decayed substantially over repeated trials. Notably, this attenuation was present selectively in neurons that responded to the stud male. In contrast, we found that initial sensory responses to single brief exposures to stud male urine as well as a range of other stimuli, remain remarkably stable following mating. Thus, we demonstrate an initially highly consistent mapping of chemosensory stimulus space in mated and unmated females that evolves over time to slowly reduce the influence of a recent significant individual, as predicted by the NTH.

Taken together, our experiments suggest a scenario that reconciles plasticity (i.e. mating-induced protection of the stud’s pregnancy blocking ability) with the essential requirement to maintain stable sensory representations that are required for guiding social and defensive behaviors. These two apparently contradictory requirements can be met because of the distinct time scales required for sensory discrimination on the one hand, and the considerably slower modulation of endocrine responses on the other.

## Results

To evaluate how mating affects representations of conspecific chemosignals, including those from the stud male, we performed extracellular ensemble recordings of AOB projection neurons while systematically testing a panel of sensory cues derived from different sexes and strains. Our stimulus set included urine from two individual BALB/c (BC) males, two individual C57BL/6 (C57) males, as well as mixed urine samples from female mice of each strain, castrated mice from each strain, and a mix of urine from different predators. We note that multiple studies have shown that exposure to an unfamiliar male urine is sufficient to induce pregnancy block, whereas urine from castrated males or female mice does not have this effect [33, 34] (but see also [35]). All recordings were made from BC females, which were either sexually naive, or mated with a male from the BC or the C57 strain (**Figure 1A**). Importantly, BC females are known to exhibit the BE with the particular male strains that we used. That is, BC females that mated with C57 males were susceptible to block following exposure to a BC male, and vice versa [36, 37]. In our experiments, females spent at least 48 hours with the stud male, and we verified mating with a vaginal plug. When a plug could not be confidently identified, mating (i.e. pregnancy) was confirmed post mortem. Recordings were made in anesthetized female mice, in a period of 5-21 days following mating, a time during which memory formation is complete and strong [38]. Recording electrodes were targeted to the external cellular layer of the AOB, which contains the cell bodies of the mitral tufted cells (MTCs, **Figure 1B**). Thus, our recordings reflect the outputs sent by AOB neurons to downstream regions. To achieve controlled and repeated stimulus delivery, we applied urine stimuli to the nostril, followed by electric stimulation of the sympathetic nerve trunk via a cuff electrode. This stimulation serves to activate the VNO and induce stimulus uptake to the vomeronasal sensory neurons that are located within the VNO (**Figure 1B**, see **Methods** and [39] for details).

**Figure 1.**
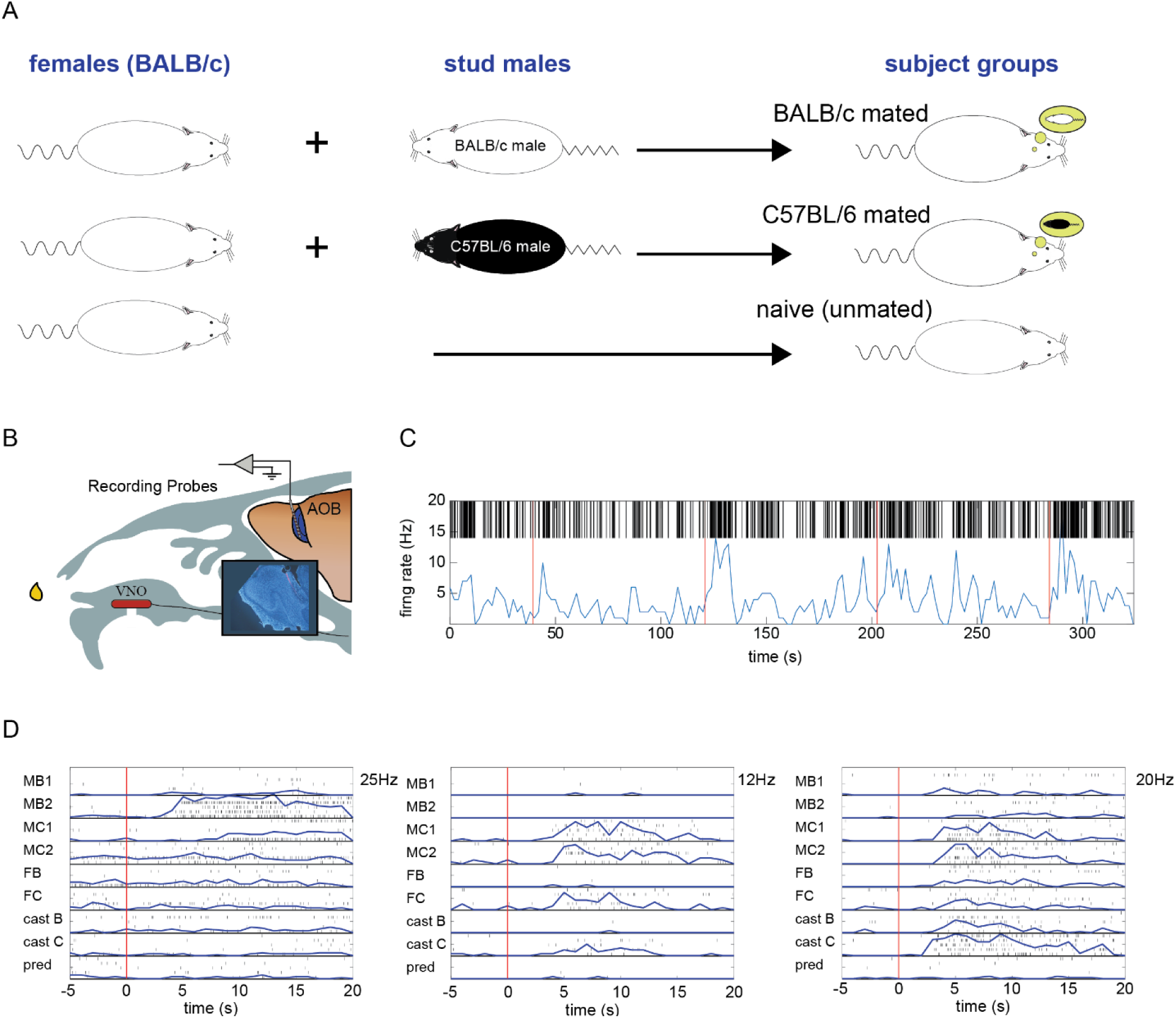
Experimental design and AOB responses to different chemosignals. **A**. Definition and icons used for the female groups in this analysis. **B**. Schematic of the recording setup. Inset shows a DAPI stained sagittal brain slice showing electrode positioning within the cellular layer of the AOB. **C**. Example of one extended trial with a spike raster and firing rates as a function of time. Red lines indicate time of sympathetic nerve trunk stimulation following stimulus application (the stud male, from the C57 strain). **D**. Examples of responses of 3 neurons (initial representations) recorded in naive females in response to the 9 stimuli. Each plot shows single trial spike times and the mean firing rates (PSTH). Red lines indicate sympathetic nerve trunk stimulation. Numbers on the upper right side of each set of responses indicate the range of the vertical scale (common to all stimulus PSTHs for each of the neurons). VNO: vomeronasal organ. AOB: Accessory olfactory bulb. Stimulus name abbreviations: MB1, MB2: two distinct male BALB/c individuals. MC1, MC2: two distinct male C57BL/6 individuals. FB, FC: urine mixes from BALB/c and C57BL/6 females. castB, castC: urine mixes from castrated BALB/C and C57BL/6 males. Pred: mix of urine from four predator species.

We used two stimulus presentation paradigms to probe AOB representations. The first is *extended* presentation, corresponding to repeated sampling of the same stimulus, and gauges how activation is maintained over time (**Figure 1C)**. The second paradigm uses interleaved single presentations of different stimuli to evaluate coding and potential memory interference, denoted here as *initial representations*. Examples of selectivity across different sexes and strains based on this approach are shown in **Figure 1D**. Interestingly, this included selectivity to individual animals of matching strain, sex and age, suggesting that the AOB could encode detailed information about individual identity in addition to broader biological characteristics.

### Selective response attenuation following extended stimulation

We begin our analysis by asking whether mating induces selective attenuation of *in vivo* responses to stud stimuli upon repeated stimulation, modeling the prolonged and repetitive investigation bout between two individuals required to elicit pregnancy block. We presented two stimuli: the stud urine and a mix of male, female, and predator urine (denoted as MFP mix, see Methods for further details). The extended stimulation paradigm (∼3 min long, see Methods for details) included four consecutive presentations without washing of the nasal cavity between consecutive stimuli (**Figure 1C**). At the end of the four repetitions, the nasal cavity and VNO were flushed with Ringer’s solution and the second stimulus in the pair was presented using the same protocol. The order of presentation of the two stimuli (stud urine or MFP mix) was randomly selected for each female.

In our analysis, we distinguish between stud responsive and unresponsive neurons, as presumably only the former were subject to plasticity following mating. Stud responsiveness was assessed based on interleaved stimulus sequences that preceded the extended stimulation protocol (described below). While stud responsive neurons could also be activated by components of the MFP mix, stud-unresponsive neurons did not show any significant response to the stud stimulus but did respond to one or more of the components of the MFP mix. In naive females, the “stud” stimulus was randomly designated using a random selection of one of the four male stimuli.

These data allowed us to evaluate the effects of learning on AOB representations by comparing sensory-driven activity in 4 groups of neurons: stud responsive and stud unresponsive neurons, in both mated and naive females (n=44 neurons from naive females, 65 neurons in mated females). The NTH model suggests some form of selective plasticity in stud responsive neurons in mated females, while *in vitro* work raised the possibility that this could take the form of time-evolving attenuation [32]. To test for such attenuation, we defined an attenuation index (AI, see methods), which measures the decrease in response magnitude from the 1^st^ to the 4^th^ presentation for each neuron. The AI essentially normalizes the firing rate of each neuron (so that neurons with higher rates do not dominate the metric) and thus assumes values from -1 to 1, with 1 corresponding to complete silencing and 0 corresponding to no change in response strength.

This analysis (**Figure 2**) supports the hypothesis that stud responsive neurons show selective attenuation in mated females upon extended stimulus presentations. The median AI values for the stud stimulus were markedly higher for responses to urine from mated relative to non-mated males (shown for all groups in **Figure 2A**). Considerably less attenuation was seen for the MFP mix, presented to the same experimental groups (**Figure 2B)**. A more complete representation of the AI distributions is shown in **Figure 2C-F**, where attenuation is not only strongest but also most significant for stud responsive neurons in mated females upon presentation of the stud urine (p=0.0033; one-tailed signrank test). Importantly, these results are also confirmed with a larger multi-unit dataset (**Figure S1**). Specifically, the shift of the AI distribution for stud responsive neurons in mated females is also most significant for the mating male (MM) stimulus. When also considering multi-unit data, a significant albeit lesser change is also observed for the MFP stimulus (**Figure S1**).

**Figure 2.**
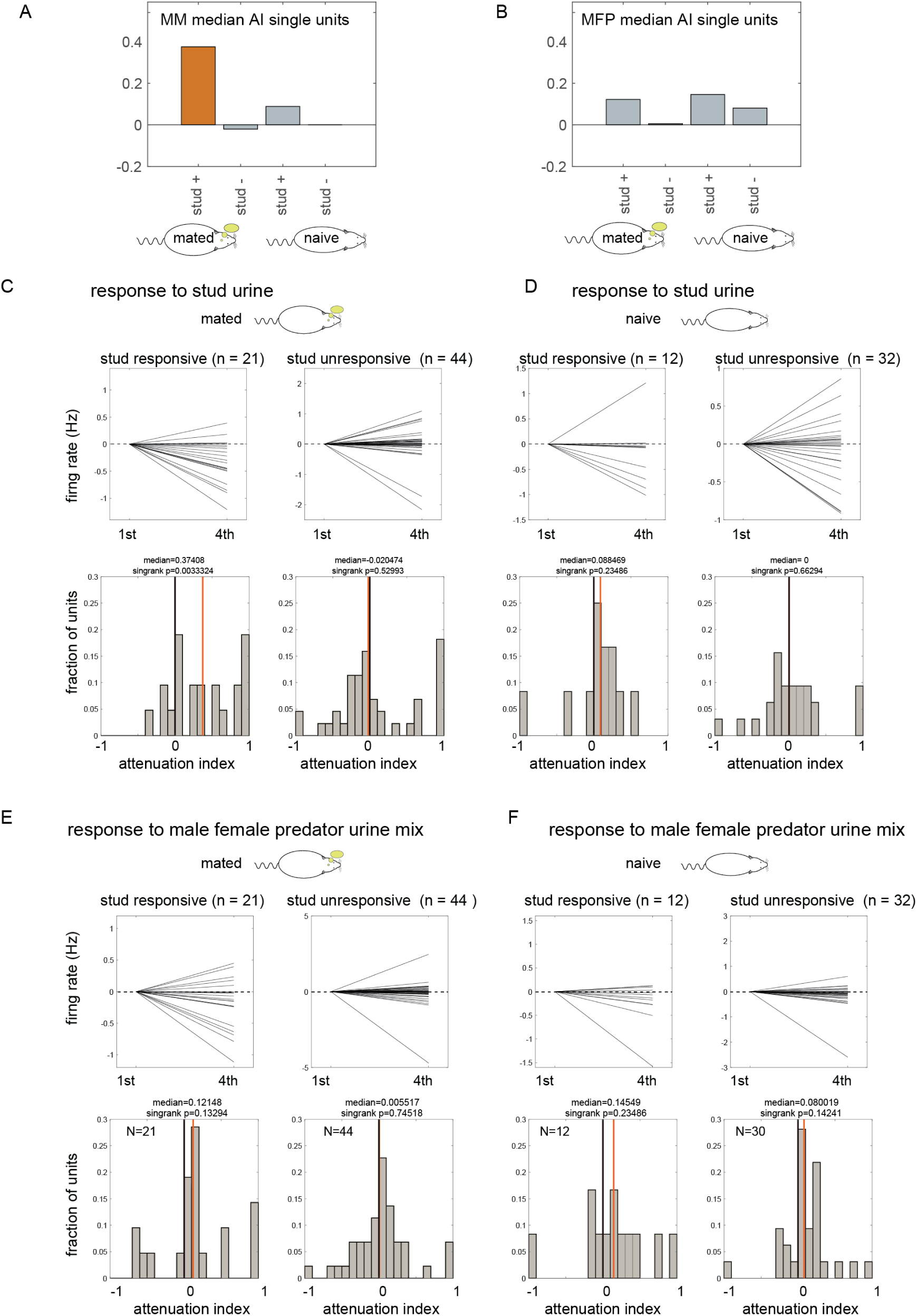
Analysis of responses to extended stimulation. **A**. Median values for the AI in response to the stud urine (MM) in each of the four groups of cells. Stud responsive neurons in mated females are indicated by the orange bar. Stud+ and stud-columns indicate stud responsive and unresponsive neurons, respectively. **B**. Median values for the AI in response to the male female predator urine mix (MFP) in each of the 4 groups. **C-F**. Detailed representation of the AI in each of the groups shown in **A** and **B. C**. Firing rates in the first and 4^th^ presentation for each of the recorded neurons in mated females in response to the stud urine, for stud responsive (left) and stud unresponsive neurons (right). Data are shown as firing rate changes relative to the first trial (which is thus 0 by definition). Lower panels: distribution of the AI for the responses shown above. In all histograms, black vertical lines indicate 0. Red vertical lines indicate the median value. **D**. Same as panel E but for neurons recorded in naive females. Here, the stud was randomly chosen as one of the four individual males. **E**. Responses in mated females to the MFP mix. **F**. Responses in naive females to the MFP mix.

Taken together, these analyses clearly indicate that mating leads to a consistent decrease in the responses of stud selective neurons upon repeated stimulation. Furthermore, attenuation appears strongest for stud stimuli and not to other stimuli that these neurons may also respond to. Overall, this analysis provides a critical *in vivo* test of how changes in firing properties measured *in vitro* translate into altered responses to biologically relevant sensory cues, and implies that the intensity with which stud responsive neurons can activate downstream pathways over an extended time frame is decreased following mating.

While we found attenuation for repeated presentations of a single stimulus, its role in coding when animals encounter more complex sequences of cues is unclear. To address this situation, we also analyzed response attenuation of these same neurons in our interleaved trials of urine from different animals (i.e. those involving *initial* representations; 5 blocks where each stimulus is presented in a pseudo random order). To test if the magnitude of the stud responses consistently decreases over the course of the ∼90-minute experiment, we calculated the AI for responses to the 1^st^ and 5^th^ presentation of stud stimuli in mated females, with responses in sexually naive females as a control where one of the males was randomly designated as the stud. For both mated and naive females, we considered only neurons that showed a significant response to the stud stimulus. The results of this analysis reveal no systematic attenuation in responses to stud stimuli in either the mated, or in the control group (**Figure S2**). Specifically, the median value of the AI was -0.18 and -0.004 for the mated and naive groups, respectively (p =0.446 and 0.308 for the mated and naive groups, respectively; one-tailed signed rank test; n=38 and n=54 for the two groups). These conclusions are further supported upon inclusion of multi-unit data, which substantially increases the sample sizes (**Figure S2**).

With these analyses, we have established that stud responsive neurons in mated females show response attenuation upon extended stimulus presentation mimicking an ongoing prolonged investigation bout. These analyses provide key experimental evidence of altered sensory responses after mating, consistent with previous *in vitro* work showing altered membrane excitability [32], and provide a potential explanation for the failure of stud odors to elicit the pregnancy block.

### Stud induced responses in naive females

The attenuation that we observe for stud-derived cues raises the possibility that this effect could interfere with the stable encoding of information about other individuals, including the activity used to recognize the stud male itself. We addressed this issue by measuring ‘initial representations’ with interleaved, single-stimulus presentations from different conspecifics. Examples of neuronal responses recorded in naive females to male stimuli are shown in **Figure 3A** (see Methods for details on the stimulation paradigm and response quantification).

**Figure 3.**
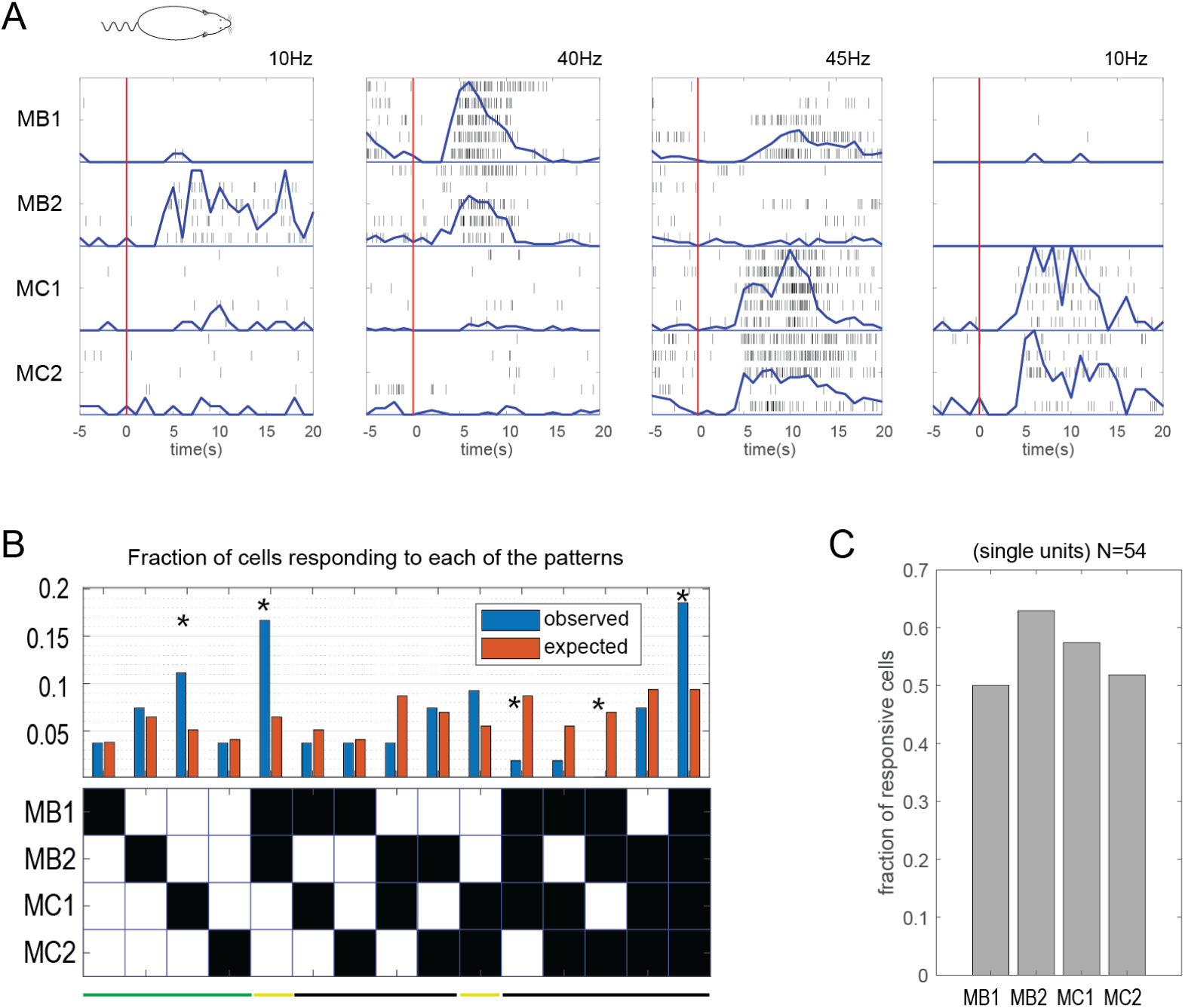
Responses to male urine in naive females. **A**. Examples of responses of four different single units recorded in naive females to the 4 male stimuli. **B**. Distribution of significant response patterns to each combination. Blue and orange bars show the observed and expected distribution, respectively to patterns indicated in the table below (a filled square in the table indicates a significant response to the stimulus shown on the left). Significant differences between the observed and expected distributions are indicated with asterisks (p<0.05, expected values calculated according to the binomial distribution, see **Table S4**). Stimulus name abbreviations are described in the legend for **Figure 1**. (N=54 single units, that respond to at least one of the male stimuli). **C**. Fraction of responsive neurons to each of the 4 stimuli across all mated females. Horizontal bars in B denote different pattern types (see text).

We first evaluated the selectivity of MTC sensory responses for the cues in our test panel (N=104 single units and 169 multi units, from 20 recording sites in 12 naive females). Overall, male urine activated a substantial portion of AOB neurons, with 39.5% and 37.5% of single units in naive female mice showing a significant response to BC and C57 urine stimuli, respectively, and 25% of the units responding to stimuli from both strains. These values remain similar when adding multiunit data (38% and 38% for the BC and C57 strains, and 23% to both strains).

We next considered how sensory responses to different categories of cues are allocated across the population of AOB neurons. Given 4 different male stimuli in our panel, there exist 15 different response patterns for each neuron defined by the set of stimuli to which it significantly responds, shown in **Figure 3B** along with the expected and actually observed number of neurons conforming to each pattern. The expected values were calculated based the marginal distributions shown in **Figure 3C** and the assumption of independence of responses to the different stimuli. Note that in this analysis we only include neurons that respond to at least one of the 4 male urine stimuli (see Table 1). Adopting the simple interpretation of the NTH, and the experiments that show that mating with an individual male from a given strain protects against pregnancy failure by another individual from the same strain [40, 41], one would expect to find neurons that show similar responses to two males from the same strain. In our data set, this includes two response patterns: BC selective neurons, and C57 selective neurons. Interestingly, both of these patterns (indicated by the yellow horizontal bars in **Figure 3B**) are more prevalent than expected by chance. Another type of response pattern, also consistent with the assumptions of the NTH, is an individual-selective neuron which would be useful for distinguishing among individuals from the same strain. Some, but not all of these patterns, indicated by green bar in **Figure 3B**, are more frequent than expected by chance. The third type of pattern includes neurons that respond to stimuli from both strains (black horizontal bar). Although these patterns generally occur less than expected by chance, together they represent a substantial fraction of AOB neurons. These latter patterns are not easily reconciled with the simplest form of the NTH, as changes in the activity of such neurons would affect representations of individuals from distinct strains, inconsistent with selective modulation of responses to stimuli from one strain. These conclusions are also supported by analyses including multi-unit data (**Figure S3, and Table S4**). Overall, we find evidence for a diverse range of response properties, including broad and relatively nonselective tuning, cells that distinguish categories such as strain, and neurons that encode finer distinctions of individual identity.

**Table 1.** Data-sets, statistics and MATLAB functions used for creating plots for each of the figures.

| Figure: | Dataset | Statistics | MATLAB functions |
| --- | --- | --- | --- |
| **Figure 2A 2B 2C 2D 2E 2F** | AOB neurons responding to mating male or one of the MFP mix components | Attenuation index as explained in the methods section. One tail Wilcoxon sign rank test | *signrank* |
| **Figure 3C** | AOB neurons (defined as neurons that showed a significant response to at least one of the stimuli) | Fraction responsive out of all cells |  |
| **Figure 3B** | Neurons that responded to at least one of the male stimuli | The percentages calculated in **3C**, were used to calculate the marginal probability of each pattern (Figure **3B**, orange). The p-value of observed vs. expected values was calculated using the cumulative binomial distribution | ***binocdf (x,n,p)*** function (with x being the observed fraction, n is the number of units, and p is the expected fraction of each pattern, calculated using the marginal probabilities of responding to each of the male stimuli in our data set. |
| **Figure 4B** | only neurons that responded to one of the male stimuli. For each strain, the maximal (in absolute value) response (out of the two responses to the two individuals) is selected. |  |  |
| **Figure 4C, 4D** | Only neurons that responded to one of the male stimuli. For each strain, the maximum response (out of the two responses to the two individuals) is selected. Only excitatory responses are used. | Explanations of the preference calculation and the stud strain suppression calculation are in given in the Methods section.<br>To test if distribution of the preferences is with median value of 0, a two-sided Wilcoxon signed rank test was used.<br>To compare if two distributions of preferences have equal median, we used a two-sided Wilcoxon rank sum test | ***signrank***<br>***ranksum*** |
| **Figure 4E** | Only neurons that responded to one of the males from the stud strain. Only excitatory responses were considered. | Explanation of the stud selective calculation is given in Methods. To test if the distribution has a median value of 0, we used two-sided Wilcoxon signed rank test. | ***signrank*** |
| **Figure 5A, 5B** | All AOB cells | To compare if two distributions of baseline firing rates (naive vs. mated) and the two distributions of lifetime sparseness has equal median, we used a two-sided Wilcoxon rank sum test | ***ranksum*** |
| **Figure 5C, 5D** | All AOB cells | correlation coefficients and the p-values for testing the hypothesis of independence between response magnitudes in mated and naive females. | ***corrcoef*** |
| **Figure 5E** | Only neurons that responded to one of the males or one of the females. Only excitatory responses were used. | Explanation of sex selective index calculation is given in Methods.<br>two-sided Wilcoxon sign rank test | ***ranksum signrank*** |
| **Figure 6A, 6B** | All AOB neurons after exclusion of units responding exclusively to castrated stimuli | Euclidean distances between responses to pairs of stimuli | ***Pdist*** |
| **Figure 6C 6D** | All AOB neurons after exclusion of units responding exclusively to castrated stimuli | correlation coefficients and the p-values for testing the hypothesis of independence between pairwise population level distances in mated and naive females. | ***corrcoef*** |

### Initial representations of the stud strain or stud individual are not *attenuated or silenced* following mating

Having characterized responses to males in naive females, we next explore how mating alters responses to male stimuli (BC mated females: N=73 single units and 120 multi units, 12 recording sites from 7 females, C57 mated females: N=49 single units and 85 multi units, 18 recording sites from 11 females). Rather than repeated presentations reflecting extended interactions with a given individual, tested above, here we considered interleaved stimuli reflecting interactions with multiple individuals where preserving accurate sensory data will be important for distinguishing between individuals. According to the NTH, responses of stud responsive neurons should be weakened following mating. However, this prediction may be interpreted in more than one way. Given that mating with one male from a given strain provides protection from another male of the same strain [40], one strong prediction is that responses to *all* males from that strain will be *silenced* following mating. Alternatively, response silencing or attenuation could apply only to neurons encoding the specific stud individual, and not to other males from the same strain.

We first test if mating leads to attenuation of responses to the stud stimuli. We distinguish between responses to the stud male strain and to the stud individual. Examples of responses of neurons recorded in mated females to male stimuli are shown in **Figure 4A**. To quantify the responses of neurons to both male stimuli from a given strain, we first characterize each neuron’s response as the larger (in absolute value) of the two responses to individuals from that strain. Inspection of the scatter plots in **Figure 4B** suggests that the relative response strength is similar for neurons from the naïve, BC-mated, and C57-mated female groups. To conduct a statistical comparison of the differences, we defined, for each neuron, a *strain preference index*. The index ranges between -1 and 1, with -1 corresponding to an exclusive response to C57 urine and 1 corresponding to an exclusive response to BC urine (See Methods). Using this index, absolute firing rates of individual neurons are normalized, yielding relative response magnitudes to the two stimuli. Statistical analysis for the naïve, BC-mated, and C57-mated female groups indicates that the median of the strain preference index is not different from 0 for any of the groups, and furthermore is not different between any pair of groups (**Figure 4C**). Group medians for each group were tested using a two-sided Wilcoxon signed rank test: p-value naive: 0.27, p-value BC mated: 0.42, p-value C57 mated: 0.99. Pairwise group comparisons were made with a two-sided Wilcoxon ranksum test: naive vs. BC mated p-value: 0.24, naive vs. C57 mated p-value: 0.75, BC mated vs. C57 mated p-value: 0.56. We then combined data from both groups, and defined a *stud strain preference index*, also ranging between -1 and 1, with 1 corresponding to an exclusive response to the stud male strain. The distribution of the stud strain preference index (**Figure 4D**) is also not different from 0 (two-tailed Wilcoxon signed rank test, p-value: 0.4885). Taken together, these analyses clearly indicate that mating does not lead to a general attenuation of initial responses to stimuli from the stud strain.

**Figure 4.**
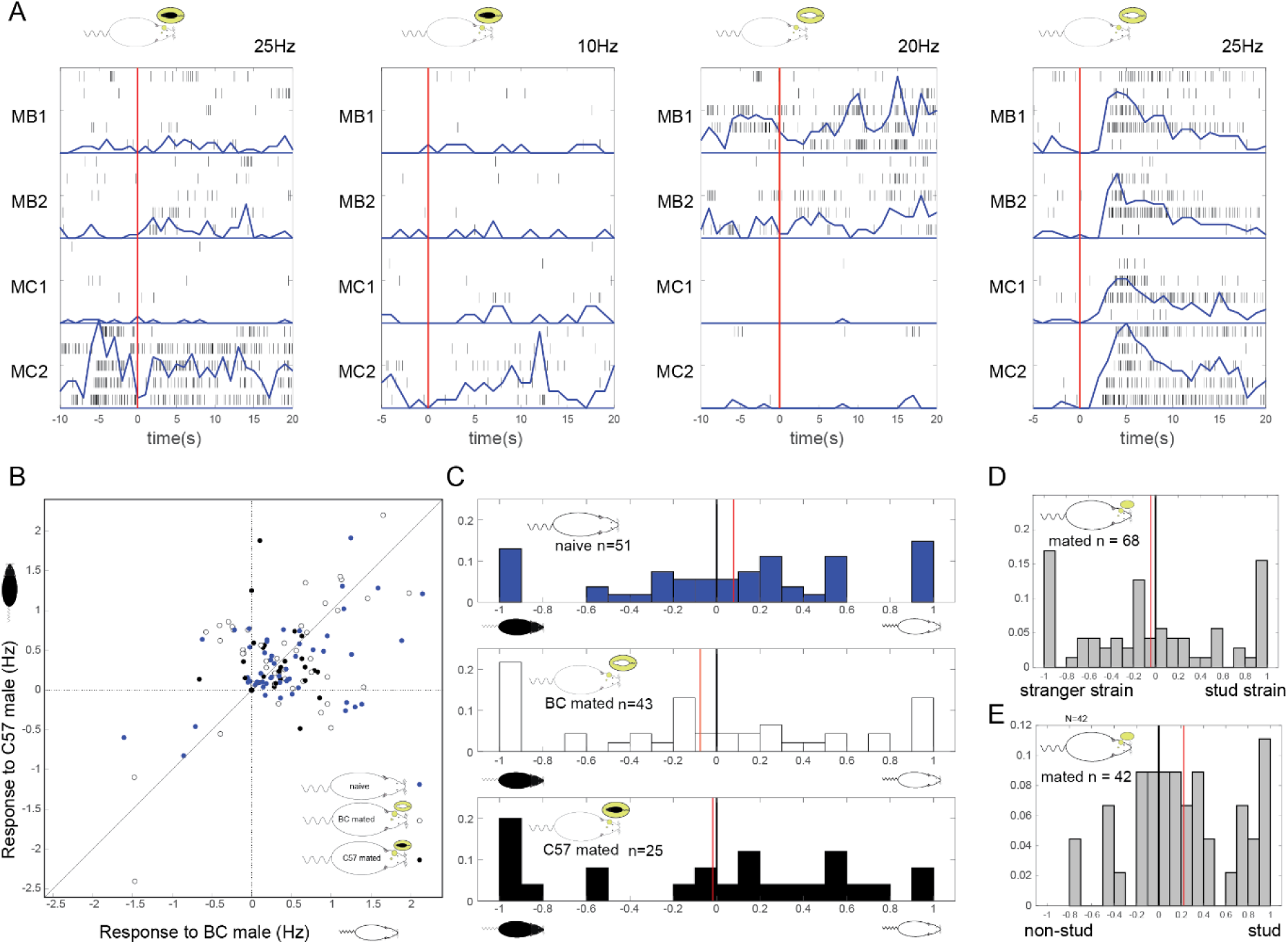
Response strength to male stimuli in naive and mated females. **A**. Examples of single unit responses to the four male stimuli in mated females (stud strains indicated by the icons next to each panel). Values above each panel indicate the maximal firing rate at the top end of the y axes. **B**. Scatter plot showing response magnitude (defined as the maximal response across both male individuals) in each of the three female groups. **C**. Strain preference indices in each of the female groups. **D**. Stud strain preference index in both mated female groups. **E**. Stud selective index in both mated female groups. Red horizontal lines in panel C, D and E indicate mean values.

Having shown no general decrease in response strength to the stud strain, we next ask if there is a selective attenuation to the stud individual. To that end, we compare the responses to the stud individual and to the non-stud individual from the same strain as the stud. Specifically, we define a *stud selective index* that ranges between -1 (reflecting an exclusive response to the non-stud individual from the stud strain) and 1 (exclusive response to the stud individual). The results of this analysis indicate that overall, responses to the stud individual are in fact somewhat stronger than responses to the non-stud individual (**Figure 4E**). The mean of the stud selective index (when neurons are pooled from both mated female groups) is 0.22, and is significantly different from zero (Wilcoxon signed rank test p=0.008).

All the analysis shown in **Figure 4** were also repeated with inclusion of multi-unit data, and the same conclusions were reached with the exception that responses to the stud individual were actually slightly stronger than to the non-stud individual (**Figure S5**). In addition, we repeated all the analysis shown in **Figure 4B-D** using the *average* response across two individuals from the same strain (rather than the *maximum*, as used above). Here too, using both single and multi-unit data, the conclusions were reached (**Figure S6**).

An alternative, complementary, approach to investigate response attenuation following mating involves the number of significantly responding neurons. A neuron’s response to a given stimulus is considered significant (see Methods) if it shows a consistent change in firing rate as compared to its baseline firing rate. According to the NTH, mating is expected to reduce the number of neurons significantly responding to males from the stud strain, or specifically to the stud individual. The results of this analysis are shown in **Figure S7**, and clearly indicate that mating does not lead to a reduction in the number of stud-individual, or stud-strain, responsive neurons.

Thus, summarizing the analyses from experiments using interleaved stimuli from a range of conspecifics, we conclude that mating does not silence or attenuate *initial* AOB MTC responses to stimuli from either the stud strain in general, or the stud individual in particular. Attenuation of sensory responses to the stud male appears to depend on extended exposure on longer timescales that are more likely to modulate hormonal state.

### Mating does not modify baseline rates, lifetime sparseness, relative response magnitude, or sex selectivity

At this stage, we can conclude that initial representations of the stud male are not changed following mating. However, mating might introduce changes in other features of AOB responses that could still affect downstream processing. To address this possibility, we first compare baseline firing rates between naive and mated females (with both mated female groups combined, irrespective of the stud strain). Specifically, we calculated mean baseline firing rate for neurons measured in each female, and compared those means across groups (**Figure 5A**). This analysis indicates that mating does not induce a consistent change in baseline firing rates across females (p=0.4127, two sample two tailed Wilcoxon rank sum test). Next, we test the selectivity of neurons in mated and naive females, using the lifetime sparseness metric (see Methods). As shown in **Figure 5B** the distribution of sparseness values across neurons does not differ for the two groups (p=0.16236 two sample two tailed Wilcoxon rank sum test), implying that mating does not lead to a general change in response selectivity.

**Figure 5.**
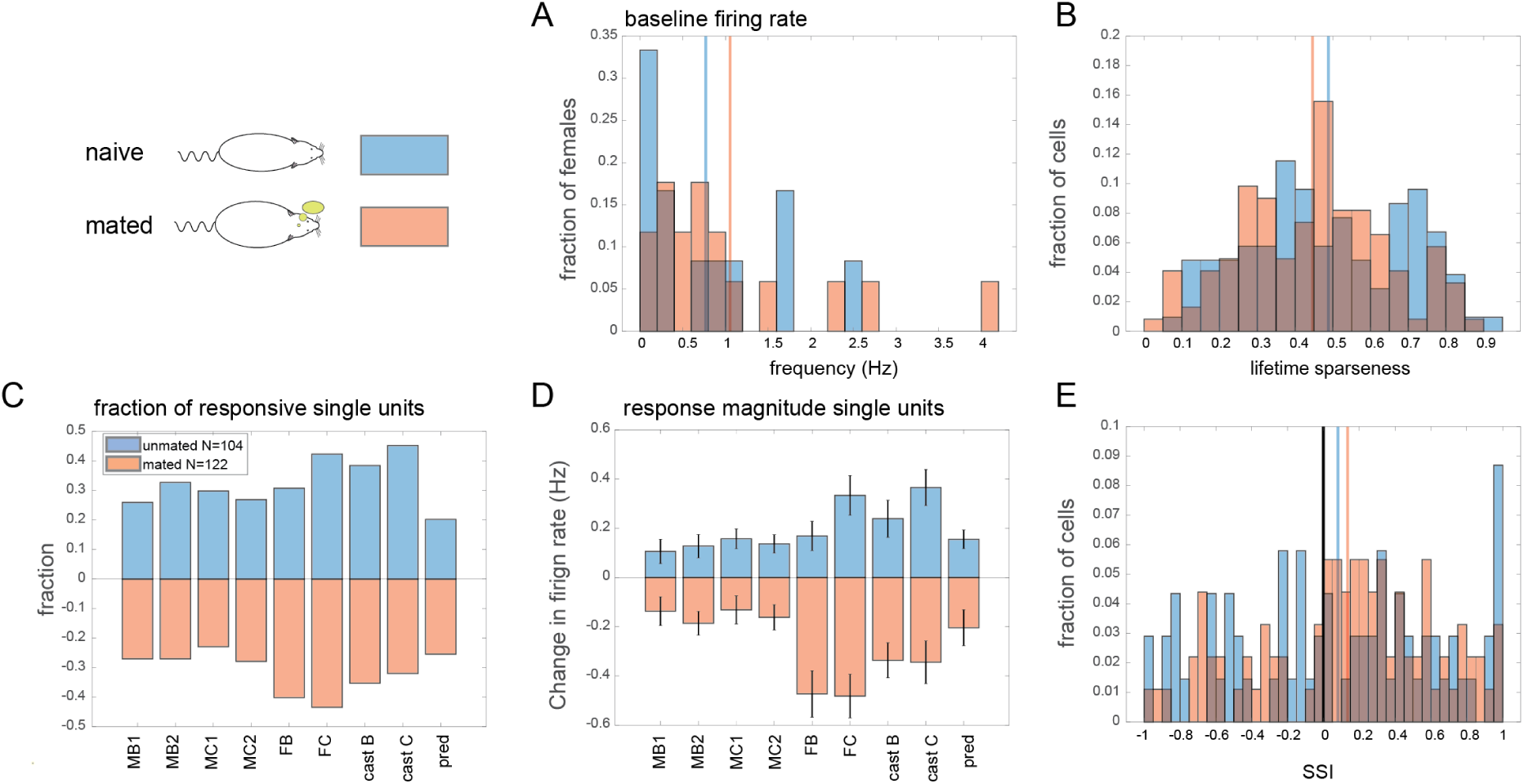
Responses to the entire panel of stimuli in naive and mated females. **A**. Distribution of baseline firing rates in naive (n=12) and mated (n=17) females. Horizontal bars represent the mean baseline firing rates. **B**. Lifetime sparseness of single neurons in the two groups. Horizontal bars represent the mean sparseness (naive: 0.4887, mated: 0.4463). **C**. Fraction of responding single neurons to each of the stimuli in the two groups. **D**. Mean (and standard errors) of firing rate changes in each of the female groups. In panels B-D: N naive=104, N mated = 122 single units. **E**. Sex selectivity index in the two female groups. Horizontal bars represent the mean SSI values (Naive: 0.0808, mated: 0.135). N naive= 69, N mated =91.

We next examined neuronal responses using our entire stimulus set which includes urine mixes from two female mouse strains, two strains of castrated males, and predator urine. Comparing the number of significant responses to each of the stimuli in naive and in mated females reveals that these values are positively correlated (**Figure 5C**, CC=0.6064 p=0.0834), and the correlation is even higher when multi-units are also included (CC=0.8 p=0.0095).

A similar analysis of the population-mean response magnitudes to each of the stimuli (**Figure 5D**), also reveals similar patterns in the mated and naive female groups, with a correlation of 0.68 (p=0.04). When including multi units, the correlation is substantially higher (0.93, p=0.0002). We note that in both groups, female stimuli elicit stronger responses than male stimuli. This is a recurring observation in AOB recordings [42-44], although in the present experiments, may also reflect the fact that female stimuli were mixes rather than individual mouse samples. Somewhat unexpectedly, castrated male stimuli mixes elicit very strong responses as well.

Despite the high and significant correlation, responses do appear to be overall stronger in mated females (**Figure 5D**), most notably for female stimuli. Statistical comparison (two-sided two sample Wilcoxon rank sum test) shows that after correcting for multiple comparisons, the only significant difference is for the female BC stimulus (**Table S8**).

Next, we asked if mating alters the *relative* response strength of male vs. female stimuli. Thus, we defined a sex selectivity index (SSI) for each neuron and compared the distribution of the index in mated and unmated females. The index ranges between -1 and 1, with a tendency to respond stronger to female stimuli reflected by positive values (see Methods). Comparison of the SSI distributions reveals no differences between the two groups (**Figure 5E**). Specifically, the mean SSI of naive female neurons is 0.0808 and not significantly different from 0 (two-sided Wilcoxon signed rank test P=0.2461). Although neurons recorded in mated females had a mean SSI of 0.135, representing a tendency to respond more strongly to females (**Figure 5E**, two-sided Wilcoxon signed rank sum test P=0.0074), the difference between the SSI distributions in mated and naive females is not significant (two-sided Wilcoxon rank sum test P=0.5933). When also including multi units, the SSI distributions assume more positive values (mean SSI naive: 0.1777, mated: 0.189) and are significantly larger than 0 in both groups (two-sided Wilcoxon signed rank test naive: p=3.9 • 10^−7^ mated: p=1.7 • 10^−10^). Yet, here too the difference between the mated and naive groups is not significant (two-sided Wilcoxon rank sum test P=0.9704). Taken together, the analyses in this section indicate that mating does not induce a global change in response strength to conspecific and predator stimuli.

### Stable population responses before and after mating

Finally, we examine if mating alters population level representation of stimulus responses. We use a Euclidean distance metric based on the firing rates elicited by each of the stimuli across all neurons, which defines a mapping between the stimuli and neuronal space where smaller distances imply greater similarity (see **Methods**). We use this metric to ask if mated and naive females share a similar mapping of stimuli at the level of the AOB. Pairwise distance matrices for single unit data are shown in **Figure 6A and B** (N= 90 and 111 single units for naive and mated females, respectively). We note that for these analyses we excluded responses to castrated males (or units that responded exclusively to castrated stimuli) as they did not show a consistent relationship with either of the other stimuli or with each other. As expected, the distance matrices show that pairs of male stimuli elicit responses that are generally similar to other male stimuli (rather than to female or predator stimuli). Then, we compared the distances for each pair of stimuli between the two female groups. As shown in **Figure 6C**, the distance correlation is highly significant, (R=0.79, P=0.000016) indicating that mating does not induce major changes in how socially relevant stimuli are mapped by AOB neurons. A more specific, direct comparison of the mapping between BC and C57 mated females shows an even higher correlation (**Figure 6D**, R=0.876, P = 0.00000018). All these conclusions remain valid when repeated using a distinct distance metric (correlation distance), when including responses to castrated males, and when multi units are included in the analysis (**Figure S9**).

**Figure 6.**
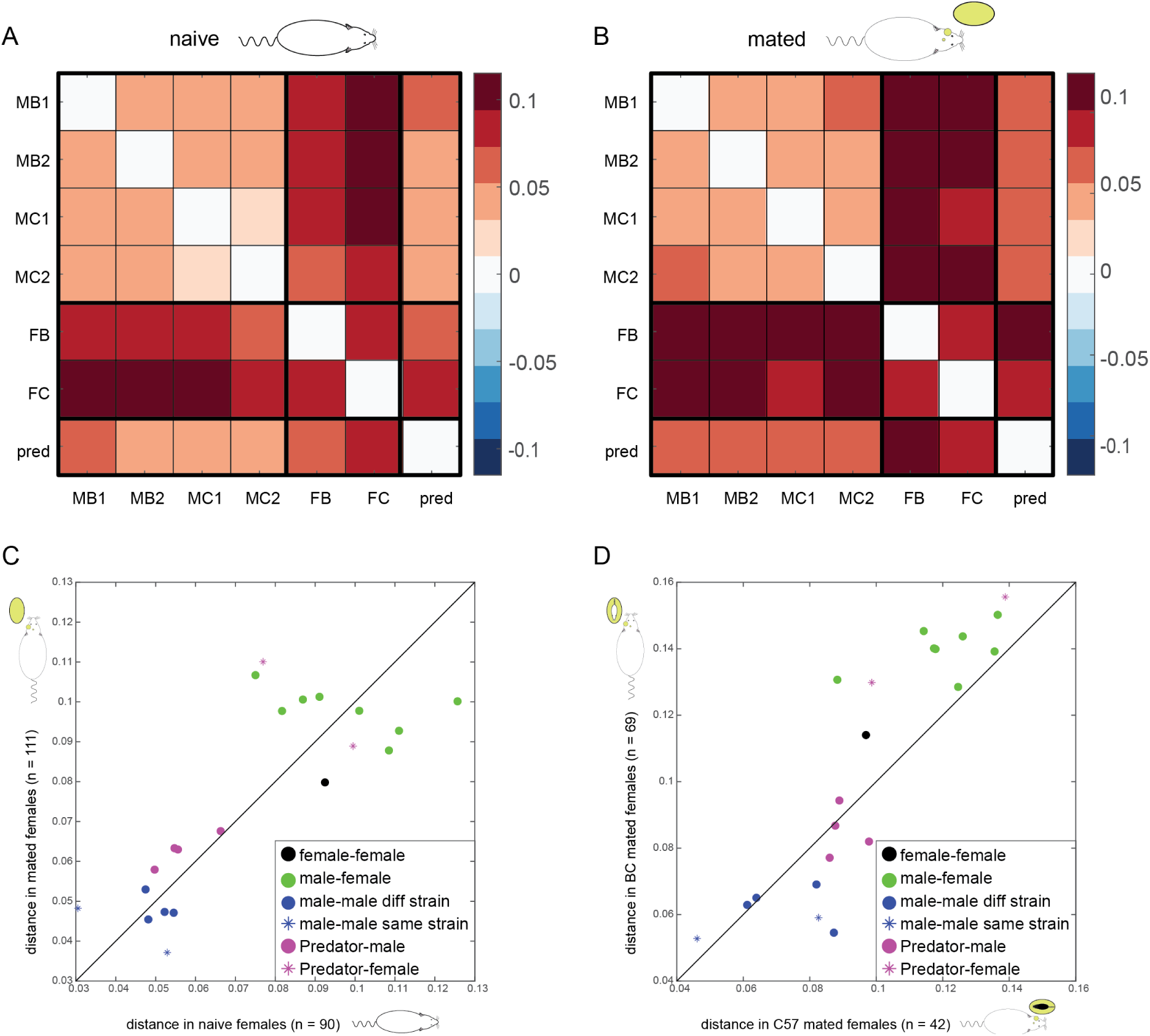
Similar population level response patterns in mated and naive females. **A**. Population response distance matrix in naive females calculated using the Euclidean distance metric. N = 90 single units. **B**. Like A, but in mated females. N = 111 single neurons. In this analysis, we only include units that respond to at least one of the non-castrated male, female or predator stimuli. **C**. Correlation of pairwise distances between mated and naive values: CC = 0.795, P=0.000016. **D**. Correlation of pairwise distances between C57 mated (n=42 single units) and BC mated females (n=69 single units): CC = 0.87, P = 0.00000018. In panels C and D, each dot corresponds to one pairwise comparison, where the markers indicate the class of the stimuli. For example, green dots indicate comparisons between a male individual stimulus and a female mix (8 different combinations).

Overall, we conclude that when the AOB receives intermingled, time-varying sensory input from multiple individuals, stimulus mapping at the level of the AOB is not altered by mating. Thus, while the attenuated response upon *repeated* stimulation with stud male chemosignals will lessen their impact on the hormonal cascade that leads to pregnancy block, *initial* representations for the first encounter with a distinct set of conspecific cues appear to remain stable on faster time scales, allowing consistent decoding of stimulus information without requiring updating of readout rules by downstream processing stages.

## Discussion

The motivation for this work was to identify the neuronal correlates of the BE in female mice. The Bruce effect is an intriguing example of behavioral imprinting of a conspecific individual that various lines of study have assigned to the AOB. Since its characterization, the dominant framework for studying the BE was the NTH, which posits that AOB responses to stud stimuli following mating would be silenced. Such silencing can readily explain the acquired immunity (with respect to pregnancy failure) of the mated female to the stud male. However, more recent data illustrating the combinatorial coding schemes of social information in the AOB [21, 29, 43, 45] highlight some potential difficulties with the NTH. Specifically, response silencing would not only render the female anosmic to her partner, but is also likely to distort representations of other stimuli, thus confounding the animal’s ability to respond appropriately to socially relevant stimuli. In this study, we conducted the first systematic *in vivo* investigation of AOB responses to conspecific stimuli in mated and unmated females. Highlighting a distinction between responses to ‘initial’ encounters with chemosignals of distinct partners, vs. prolonged exposure to cues from the same stud male, our results reconcile the ability to form a memory of the stud male that will block his ability to induce pregnancy block, while otherwise preserving a stable encoding scheme that obviates the need to modify decoding rules readout by downstream processing stages. The disassociation appears to exploit the distinct time scales of behavioral interactions and endocrine processes. While the VNS is inherently a slow system, behavioral interactions such as mating and fighting that require identification of features such as sex, genetic relatedness, and familiarity, can be initiated by relatively rapid sensory sampling that can occur within seconds. In contrast, time scales associated with endocrine processes are generally longer, on the order of minutes to hours. Here, we show that while initial stimulus presentations remain highly stable following mating, prolonged stimulation is associated with a selective attenuation of responses to the stud male.

The NTH gained popularity due to its elegance, the multiple converging lines of evidence that supported it [46-48] [22, 49], and the clear predictions it raised concerning how AOB representations of the stud male should change following mating. Specifically, the NTH posits that mating will attenuate stud responsive neurons [3]. According to this idea, stud male stimuli would no longer activate AOB mitral cells, and hence could not trigger the cascade of events that would otherwise induce pregnancy block. Along with its elegance however, the NTH raised two conceptual difficulties. First, a scenario in which a female becomes anosmic to her mate, even if only via the VNS, seems non-adaptive. The second difficulty is associated with crosstalk – namely the observation that most AOB units are activated by a broad range of stimuli, so that neuronal activity associated with specific stimuli involves overlapping neuronal ensembles. When the NTH was first proposed, there were no direct recordings of AOB activity, and while the first single neuron recordings from behaving mice supported the idea that individuals are sparsely represented by the activity of AOB projection neurons [20, 50], subsequent studies painted a more complex picture, involving a distributed code [25-30]. Furthermore, it is now clear that the list of “pregnancy blocking” elements is not confined to one or few classes of stimuli. Proposed triggers include low molecular weight constituents of urine [37], MHC peptides [36], ESP peptides [51], and mitochondrial encoded maternal peptides [35]. This observation underscores the assumption that a substantial proportion of AOB neurons is involved in individual recognition, further arguing against a narrow labelled line code in which individuals are represented by the activity of a highly selective, dedicated, group of neurons. Moreover, a large fraction of AOB neurons respond to stimuli associated with highly distinct behavioral outcomes [21]. Thus, any mechanism that involves silencing of activity of any subset of neurons, is likely to modify representations of other stimuli as well. Yet, our observations suggest that mating does not modify initial stimulus representations, circumventing both of the potential problems raised here.

It should be noted however, that in principle, the same issues of cross talk may also apply to extended stimulation. This could present a problem if stud responsive neurons are also activated by other male individuals, as is often the case, and implies that mating with one male could protect from the pregnancy blocking effects of other males. The extent of “protection” will depend on the degree of overlap between neuronal populations responding to distinct individuals or stimuli. This is consistent with the observation that mating with an inbred male from a given strain, will protect from pregnancy block from another individual of the same strain [40], even though different same-strain individuals do not elicit identical neuronal responses (**Figures 3, 4**). However, unlike the process of decoding the fine features associated with a given sensory stimulus, which likely involves a combinatorial code across multiple neurons [29], the effect on endocrine processes is more likely to follow a mass-action rule. Thus, despite a certain degree of overlap with stud responding neurons, the neuronal population activated by stranger males’ stimuli does not likely undergo sufficient attenuation (upon prolonged stimulation) to cancel their pregnancy blocking effect.

Mechanistically, a key tenet of the NTH, and other models of learning in the olfactory bulb, is that learning involves the coincidence of stud responsive neurons with noradrenaline (NA) and that this in turns leads to selective increases in inhibition of MTCs by granule cells. Various groups have studied the effects of NA on AOB physiology. Conceptually, the most relevant study examined the effects of NA in the MOB [52] and it showed selective silencing of responses to stimuli after concomitant presentation with NA administration. These results are similar to those expected in the AOB following mating, but involve a shorter time scale than is relevant for the Bruce effect. Other studies examined the effects of NA on AOB responses in brain slices, or ex-vivo AOB presentation, the latter of which enable stimulus delivery and thus measurement of the effects of NA on stimulus responses. Studies of the (short term) effects of NA on AOB slices yield a complex picture. Some studies have shown evidence for disinhibition of MTC activity [53], while others suggest the opposite, namely that increased GABAergic signaling gives rise to silencing of AOB MTC activity [54, 55]. A more recent study using an ex-vivo preparation showed an overall mild decrease in MTC responses, but the effect was quite heterogeneous, both across neurons and across stimuli for a given neuron [56]. These studies involve shorter time scales than that examined here, and paint a complex picture that likely reflects the conditions that facilitate memory formation rather than the long-term storage of information. It may well be that experience induced changes in response to extended stimulation involves modification of either MTC or granule cell excitability, as well as changes in the synaptic strength of MTCs and granule cells [31] [57]. We note that our findings suggest that the same stud responsive neurons show stronger attenuation for the stud male stimulus than for the MFP urine mix (**Figure 2**). Such stimulus specificity is not easily explained solely by changes in MTCs excitability, which would likely affect all activating stimuli similarly.

Although our observations help reconcile memory formation with maintained sensory range and acuity, we cannot confirm that the observed attenuation is sufficient to confer resistance to the stud male, nor can we entirely exclude the possibility that mating may alter initial representations under other conditions. For example, our experiments were conducted under anesthesia. The proposed mechanism according to the NTH is increased inhibition upon stud selective MTCs [3], and one may argue that under anesthesia the effects of inhibition are diminished [58-60]. However, it has been shown that inhibition substantially shapes AOB responses in anesthetized mice [42], and unlike the main olfactory bulb (MOB), no prominent differences in baseline activity and response properties were reported between the awake and anesthetized AOB [21, 42, 50]. Yet, even if inhibition is reduced in the anesthetized preparation, one would expect to see at least some degree of attenuation, and this is inconsistent with our observations. A related possibility is that learning involves changes in the pattern of feedback projections to the AOB [61-63], and the effects of feedback projections may be weakened during anesthesia. For example, one could argue that imprinting involves selective feedback inhibition of stud related MTCs, and if the feedback is entirely silenced during anesthesia, then our experiments would not reveal the physiological correlates of learning. While this possibility cannot be entirely refuted, it not only depends on the unlikely assumption of complete lack of feedback activity during anesthesia, but also assigns the representation of the individual and its modification following mating to downstream projection areas, an idea which is inconsistent with our current understanding of VNS function.

To summarize, in the context of learning in the Bruce effect, our present work reconciles memory for the stud male with maintaining constancy of stimulus representations in the AOB. These apparently contrasting ends can be achieved via the different temporal scales required for sensory processing to guide behavior (corresponding to initial representations) on the one hand, and those required for affecting downstream endocrine processing on the other. Thus, our results are consistent with the NTH, but in a manner that is distinct from its original formulation. Future studies should directly test how extended AOB stimulation in response to learned and non-learned stimuli, affects responses in downstream processing stages of the VNS, and the mechanisms that underlie this attenuation.

In broad terms, our study highlights a form of multiplexing, a scenario in which the same channels convey distinct dimensions of information in parallel. In the present case, information about chemosensory stimuli serves to guide both ongoing behavioral interactions, and on a different time scale, slower acting endocrine processes. This temporal separation allows changes to take place in one temporal scale with minimal interference to the other and could serve in other contexts and sensory systems. Indeed, virtually all sensory systems detect and convey signals at various temporal scales that correspond to distinct stimuli and may give rise to specific outputs. It remains to be explored to what extent the phenomenon observed here applies in other contexts and sensory systems.

## Methods

### Mice

All procedures were approved by the ethical committee of the Hebrew University Medical School. The dataset includes recording from 12 naive (unmated) and 18 mated BALB/c females. Stimuli were collected from adult male and female mice of the BALB/c and C57BL/6 strains. All mice were purchased from Envigo Laboratories (Israel).

### Stimuli

For urine collection, mice were gently held over a plastic sheet until they urinated. The urine was transferred to a plastic tube with a micropipette and then flash frozen in liquid nitrogen and subsequently stored at −80°C. For stimulation, urine was diluted in Ringer’s solution (1/10). To reduce the effects of sample-to-sample variation in individuals’ urine, stimuli were pooled samples collected from at least 3 different days. Each female was exposed to 9 different stimuli as listed in **Figure 1**. Castrated male urine was collected from 8 weeks old males that were castrated (testis removal under Isoflurane anesthesia) at the age of 4 weeks (before puberty). Castrated male urine and the female urine mix included a mix of at least 3 individuals from each strain (in the case of females, the estrus stage was not controlled). Predator urine included a mix of four species of predator urine (bobcat, lion, fox and wolf, purchased from PredatorPee (Maine, US).

### Experimental design

The experimental procedures are described in detail in [39]. Briefly, mice were anesthetized, a tracheotomy was performed with a polyethylene tube, and a cuff electrode was placed around the sympathetic nerve trunk. Incisions were closed with veterinary glue, after which the mouse was placed in a custom-built stereotaxic apparatus. A craniotomy was made, the dura was removed around the penetration site, and electrodes were advanced into the AOB at an angle of ∼30° with a manual micromanipulator (model MM-33, Sutter Instruments). All recordings were made with 32-channel probes (A4×8-5 mm-100-200-177-A32 or A4×8-5 mm-50-200-177-A32 configurations; NeuroNexus, Michigan, US). Before recordings, electrodes were dipped in fluorescent dye (DiI, Invitrogen) to allow subsequent confirmation of electrode placement within the AOB external cell layer that contains MTC bodies [64]. In the beginning of every experimental session, we recorded ∼10 minutes of neuronal activity, with no stimulus presentation (spontaneous activity). This period was followed by recording of neuronal activity during stimuli presentation.

### Stimulus delivery

In each trial, 2μl of the stimulus were applied directly into the nostril (stimulus application). This application is given manually by the experimenter, upon prompting by a sound sequence generated by the session management program (custom-written in MATLAB, see below). After a delay of 20 s, a square-wave stimulation train (duration, 1.6 s; current, ±120 μA; frequency, 30 Hz) was delivered through the sympathetic nerve trunk (SNT) cuff electrode to induce VNO pumping and stimulus entry to the VNO lumen (SNT stimulation). Throughout the manuscript, we refer to the start of the stimulation train as time 0. Following another delay of 40 s, the nasal cavity and the VNO were flushed with 1–2 ml of Ringer’s solution, which flowed from the nostril into the nasal cavity and drained via the nasopalatine duct using a solenoid-controlled suction tube. The flushing procedure was 40s long and included a single sympathetic trunk stimulation to facilitate stimulus elimination from the VNO lumen. In each session, the 9 different stimuli were presented in a pseudorandom order, typically five times each. In a subset of experiments, an additional block of repetitive exposure was included at the end of the session. These blocks included only 2 stimuli. The stud male urine and a mix of urine from males, females and predators (denoted as MFP mix). The males in the MFP mix were the two individuals from the non-stud strain, while the females were from both strains. In naive females the “stud” was randomly selected out of the 4 males. Each stimulus was presented in 4 consecutive trials. Each trial was composed of a 10 s ITI, stimuli application to the nostril, 20 s delay, stimulation of the SNT via the cuff electrode and additional 40 s delay. Importantly, in these sessions, the stimulus was not washed after each delivery. At the end of the 4 repetitions, the nasal 0. cavity and VNO were flushed with 1–2 ml of Ringer’s solution. This subset of experiments included recordings from 3 unmated females and 5 mated females (one mated with BC male and 4 mated with C57 males).

### Hardware and experimental control codes

The experiments were controlled using custom-written MATLAB code. The MATLAB code determined the pseudo-random order of stimuli, and interacted with the data acquisition card (I/O board, e.g., a National Instruments USB 6343 board) which controlled the speaker, a solenoid valve and the electrical stimulus isolator (AM-systems model 2200). Neuronal data were recorded using an INTAN board (RHD2000 V1, Intan Technologies).

### Data processing

Signals were sampled at 25 kHz and bandpass filtered (300–5000 Hz). Spike waveforms were extracted using custom-written MATLAB code. Spikes were sorted automatically using the KlustaKwik program [65] and then manually verified and adjusted using the Klusters software [66]. Spike clusters were evaluated by their spike shapes, projection on principal component space (calculated for each session individually), and autocorrelation functions. A cluster was designated as a single unit if it showed a distinct spike shape, was fully separable from both the origin (noise) and other clusters along at least one principal component projection, and if the inter-spike interval histogram demonstrated a clear trough around time 0 of at least 10 ms. Clusters not meeting these criteria were designated as *multi-units*. Multi-units are assumed to be composed of a mix of a small number of units with small signal amplitudes that do not allow confident separation according to their unique spike form.

#### Selection of units for analysis

Unless indicated otherwise, we considered all neurons that were identified as AOB neurons (see Table 1). Specifically, we included neurons that showed significant stimulation-locked responses to at least one of the tested stimuli. A stimulation-locked response is considered significant if the distribution of single-trial firing rates (typically, five single-trial values for each stimulus), quantified for 40 s following VNO stimulation is significantly different from the distribution of the pre-stimulus firing rates of the same neuron. The pre-stimulus firing rate distribution was evaluated during the 15 s period before stimulus application, pooled across all trials of all stimuli for each neuron. The response of a neuron to a given stimulus is considered significant if the set of firing rate responses following the 5 repetitions of odor exposures differ at the *p* < 0.05 significance level from pre-stimulus firing rate, determined using ANOVA (*anova1* function in MATLAB). Once included, all individual trials for all stimuli from that neuron were used for the analysis.

### Data analysis

All data analyses and visualizations were performed with custom-written MATLAB programs. Response magnitude (ΔR) is defined as the difference between firing rate following SNT activation and the baseline firing rate during the 15 s inter trial interval (ITI). For the repetitive exposure experiment which did not include flushing of the nasal cavity between consecutive stimulations, the activity before a given stimulation may be influenced by the previous presentation. Hence the analysis of this set of experiments is not based on response magnitude as defined above, but rather on the actual firing rate in the 40 sec post-stimulation period.

#### Preference indices

In order to avoid inter-neuron variability and to compare responses to two different stimuli, we applied preference indices of the form:

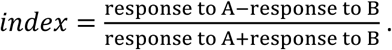

Where A and B are two stimuli under comparison. In this calculation, the preference of each neuron is normalized to values between -1 and 1, where 1 indicates an exclusive response to A, and -1 indicates an exclusive response to B. A value of 0 indicates equal responses to both stimuli. Because the index is only meaningful for positive responses, it was only computed for the set of cells that showed a significant excitatory response to one or more of the stimuli under comparison, where all responses with rate decreases were set to zero.

#### Strain preference index

For the strain preference index, A = max ΔR BC (the larger of the responses to the two BC males), and B = max ΔR C57 (the larger of the two responses to the two C57 males). For this analysis we only included neurons that responded to one of the 4 male stimuli.

#### Stud strain selective index

For the stud selective index, A = max ΔR stud (the larger of the responses to the two stud strain males), B= max ΔR unfamiliar male (the larger of the two responses to the two unfamiliar strain males). For this analysis we only included neurons that responded to one of the 4 male stimuli.

#### Stud selective index

For the strain preference index, A = ΔR to the stud male individual, B = ΔR to the non-stud individual. For this analysis we included neurons that responded to at least one of the two males from the stud strain

#### Sex selectivity index (SSI)

For the SSI, A = mean ΔR to male stimuli, B= mean ΔR to female stimuli. For this analysis we only included neurons that responded to one of the four males or one of the two female stimuli.

#### Attenuation index

For the attenuation index, A is the firing rate during the 40 s period after the first exposure, while B is the firing rate following the fourth exposure. For this analysis, we only included neurons that responded to one of male, female, or predator stimuli.

#### Sparseness

Lifetime sparseness is a measure that quantifies selectivity of individual neurons based on a given stimulus set. Lifetime sparseness (S) was computed using the definition given in [67]:

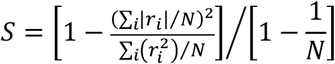

Where *r*_*i*_ is the response of neuron to the i’th stimuli (averaged across all trials) and N is the number of stimuli (N=9). S varies between 0 and 1, such that 0 indicates uniform responses to all stimuli whereas 1 indicates a response to only one stimulus. Sparseness was calculated for the data-set of single units.

#### Measures of population distance

Distances between response vectors were calculated using both the *Euclidean* distance and the *correlation* distance (defined as 1 minus the correlation between responses to different stimuli). Distances were calculated to all stimulus pairs (7 stimuli, corresponding to 21 unique pairs, as responses to castrated male urine were not included in this analysis). Pairwise distances between response vectors were calculated using MATLAB *pdist* function using the Euclidean and correlation distances. The correlation between the pairwise distances was calculated for the response magnitude using the *corrcoef* function in MATLAB.

## Supplementary files

**Figure S1**. Attenuation index including single and multi-units. Related to Figure 2.

**Figure S2**. Attenuation index for responses to the 1^st^ and 5^th^ presentation of stud stimuli. Related to Figure 2.

**Figure S3**. distribution of response patterns to male stimuli in naive females using single and multi-unit data. Related to figure 3.

**Table S4**. Statistical analysis of response pattern frequency under the binomial distribution. Related to figure 3.

**Figure S5 (parts 1-3)**. Analysis of response strength to male stimuli from both male strains using both single unit and multi-unit data. Related to Figure 4.

**Figure S6 (parts 1-3)**. Analysis of response strength to male stimuli from both male strains when response strength is characterized as the average rather than the maximal response. Related to Figure 4.

**Figure S7**. Distribution of significant responses to male stimuli in naive and mated females.

**Table S8**. Comparison of median response magnitudes between naive and mated females. Related to Figure 5.

**Figure S9 (part 1-3)**. Comparison of pairwise population level distances using different variations. Related to Figure 6.

## Acknowledgements

This research was funded by a BSF grant (3013002171) to YBS, IGD and SDS. The authors thank anonymous members of the PhD committee of M.Y.F for providing insightful comments about this work. We thank Mrs. Oksana Cohen for castrated urine stimuli.

