## Supplemental figures and tables for "Balancing memory fidelity and representational stability in the female mouse accessory olfactory bulb"

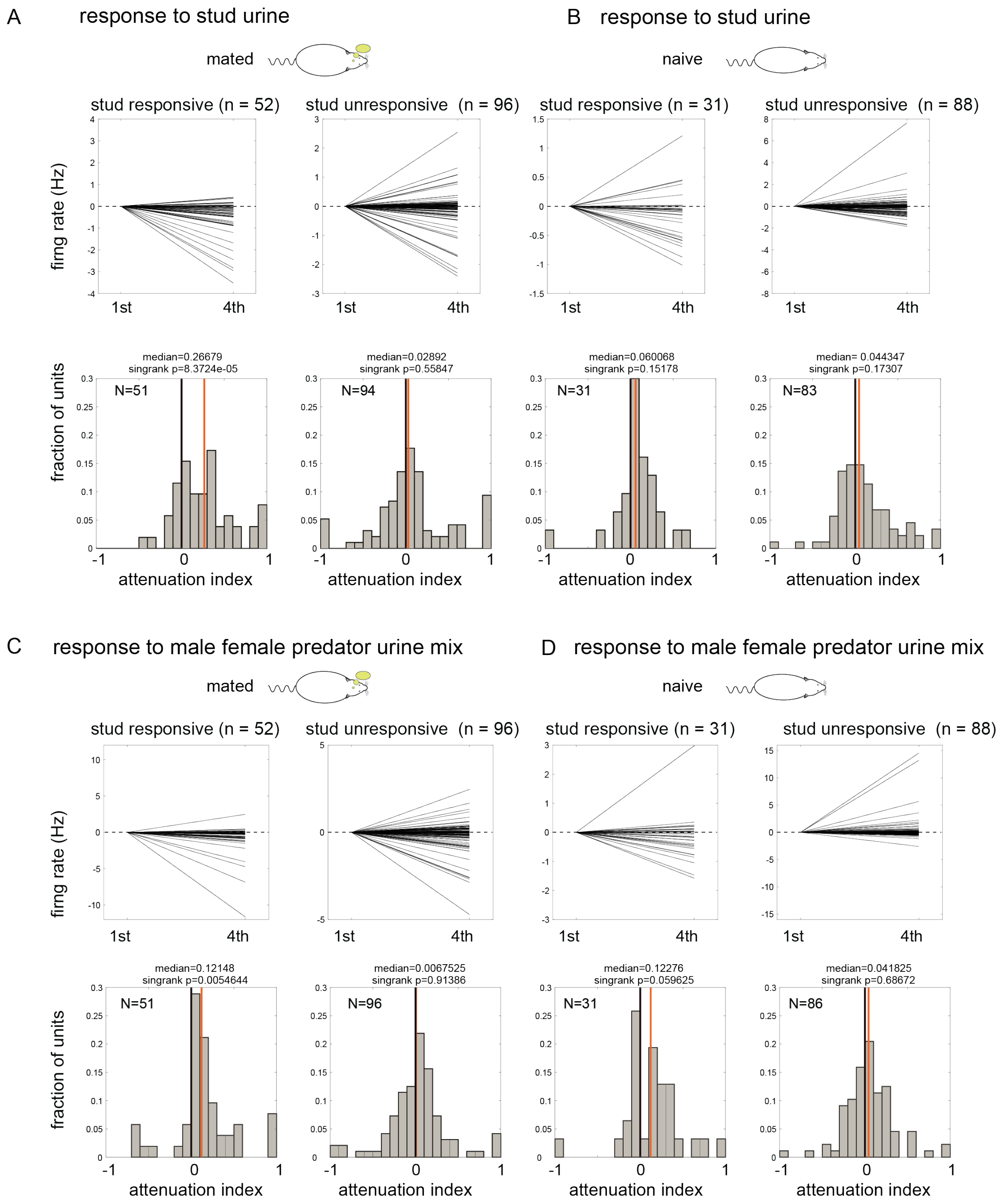


**Figure S1. Attenuation index including single and multi units**. Same format as Figure 2 (panels C, D, E, F).

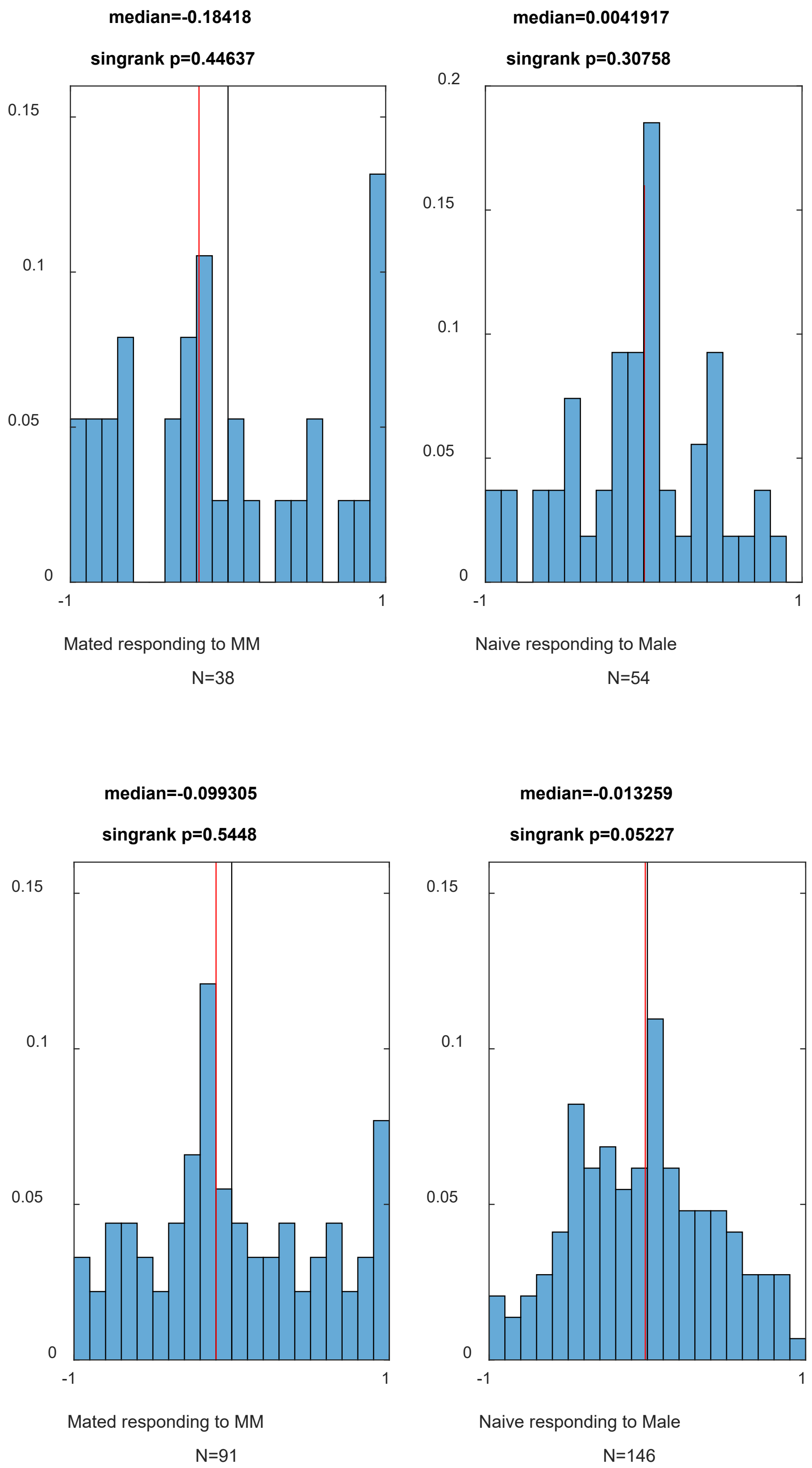


**Figure S2. Attenuation index for responses to the 1st and 5th presentation of stud stimuli**. For both mated and naive females. For mated females, we considered neurons that showed a significant response to the stud stimulus. For naive females, we considered neurons that respond significantly to at least one of the males. The results of this analysis reveal no systematic attenuation in responses to stud stimuli in either the mated, or in the control group. Top panels show AI distributions for single units recorded in mated (left) and naive (right) females. Bottom panels show AI distributions in the same groups upon inclusion of multi units as well.

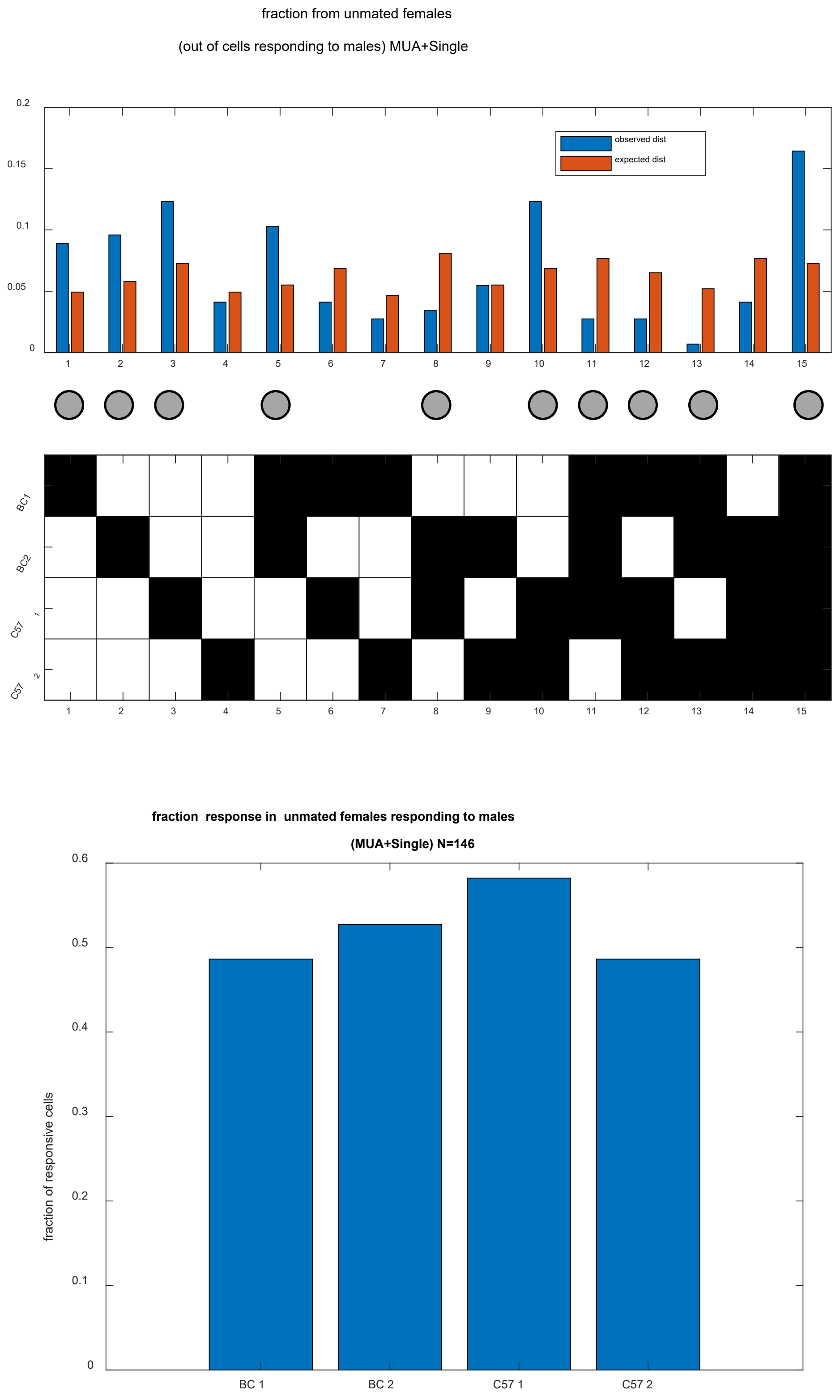


**Figure S3. distribution of response patterns to male stimuli in naive females using single and multi unit data.** Top panel is analogous to Figure 3B and bottom panel is analogous to 3C, but here multi unit data is also include. BC1, BC2, C571, C572 correspond to MB1, MB2, MC1, and MC2 used in the main text. Significant deviations from the expected frequencies are indicated by gray circles (see **Table S4** for exact values). Note that the trends observed with single unit data (Figure 3) are also observed here with multiunit data.

| | P exp bigger than obs | P exp smaller than obs (1-P) |
|---|---|---|
| 1 | 0.66 | 0.34 |
| 2 | 0.73 | 0.27 |
| 3 | 0.98 | 0.02 |
| 4 | 0.62 | 0.38 |
| 5 | 0.998 | 0.002 |
| 6 | 0.47 | 0.53 |
| 7 | 0.62 | 0.38 |
| 8 | 0.14 | 0.86 |
| 9 | 0.68 | 0.32 |
| 10 | 0.92 | 0.08 |
| 11 | 0.045 | 0.95 |
| 12 | 0.19 | 0.80 |
| 13 | 0.02 | 0.98 |
| 14 | 0.42 | 0.58 |
| 15 | 0.99 | 0.01 |

| | P exp bigger than obs | P exp smaller than obs (1-P) |
|---|---|---|
| 1 | 0.9864 | 0.0136 |
| 2 | 0.9766 | 0.0234 |
| 3 | 0.9903 | 0.0097 |
| 4 | 0.4157 | 0.5843 |
| 5 | 0.9932 | 0.0068 |
| 6 | 0.1191 | 0.8809 |
| 7 | 0.1837 | 0.8163 |
| 8 | 0.0188 | 0.9812 |
| 9 | 0.5873 | 0.4127 |
| 10 | 0.9945 | 0.0055 |
| 11 | 0.0108 | 0.9892 |
| 12 | 0.0358 | 0.9642 |
| 13 | 0.0036 | 0.9964 |
| 14 | 0.0633 | 0.9367 |
| 15 | 0.9999 | 0.0001 |

**Table S4. Statistical analysis of response pattern frequency under the binomial distribution.** Top: single unit data. Bottom: Single and multi-unit data. The left column indicates the pattern number (corresponding to the columns shown in Figure 3B for single unit data, and Figure S3 for single and multi unit data). The center and right columns indicate the probability to a number larger than, or smaller than, the observed values under the null hypothesis of independence between responses to distinct stimuli. Note that patterns that are significant with single units are also significant with multi units included.

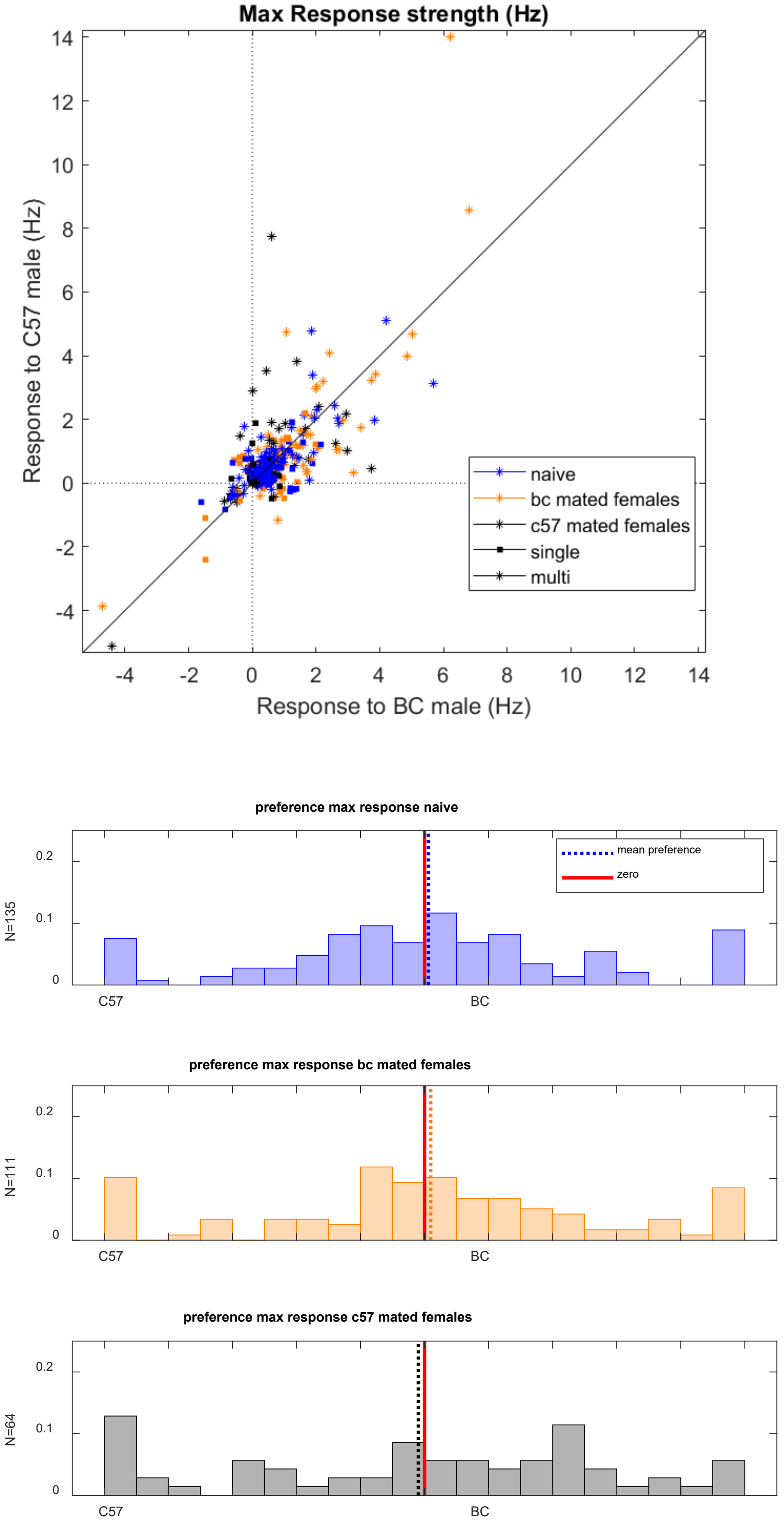


**Figure S5 (part 1). Analysis of response strength to male stimuli from both male strains using both single unit and multi unit data.** With the exception of some formatting changes, the top and bottom panels are entirely analogous to those shown in **Figure 4B** and **Figure 4C**, respectively.

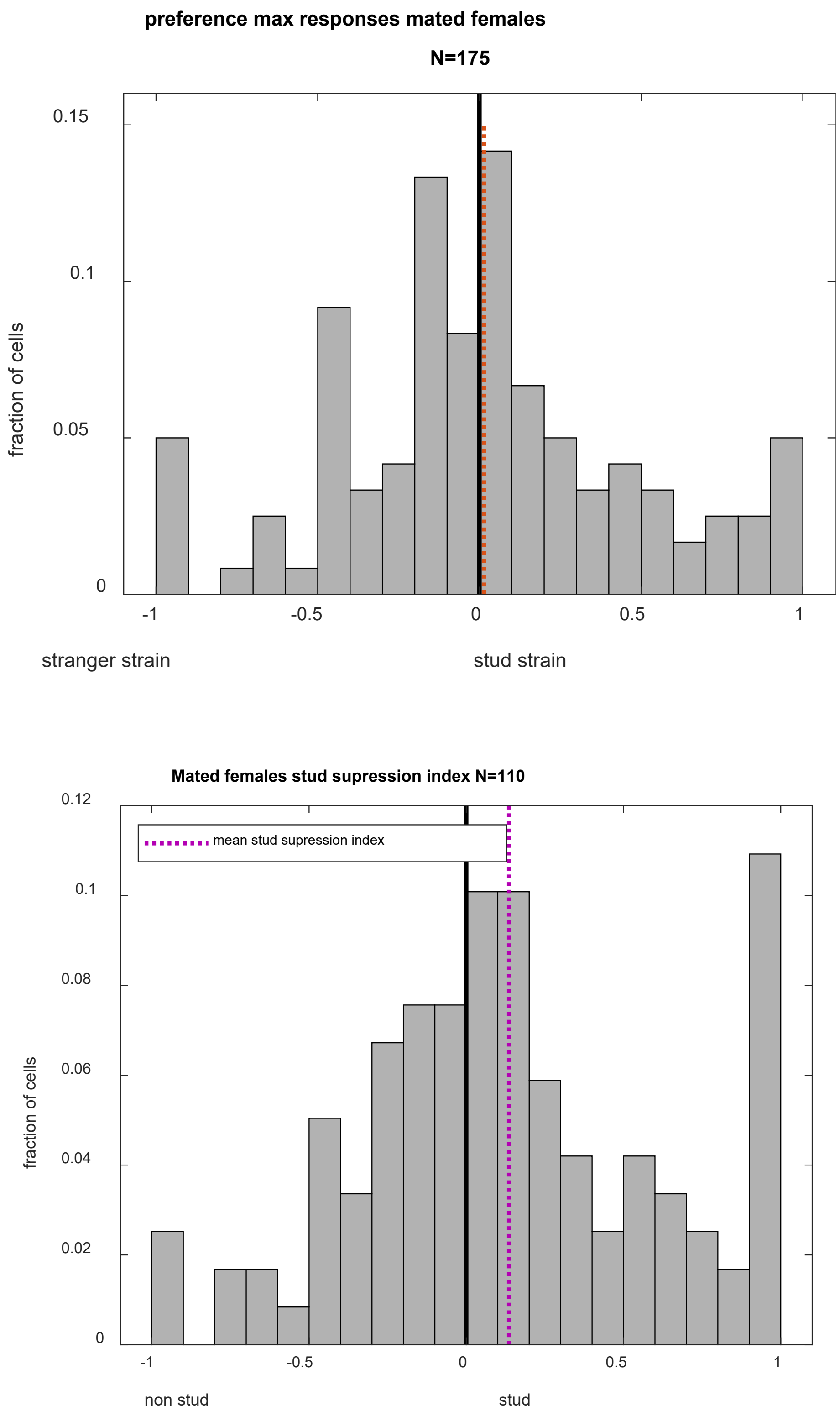


**Figure S5 (part 2). Analysis of response strength to the stud strain and stud individual using single and multi unit data.** With the exception of some formatting changes, the top and bottom panels are entirely analogous to those shown in **Figure 4D** (top panel here) and **Figure 4E** (bottom panel here), respectively. Top: mean index: 0.0192 , pval (sign rank test): 0.7701. Bottom: mean index: 0.1358, pval (sign rank test): 0.0142

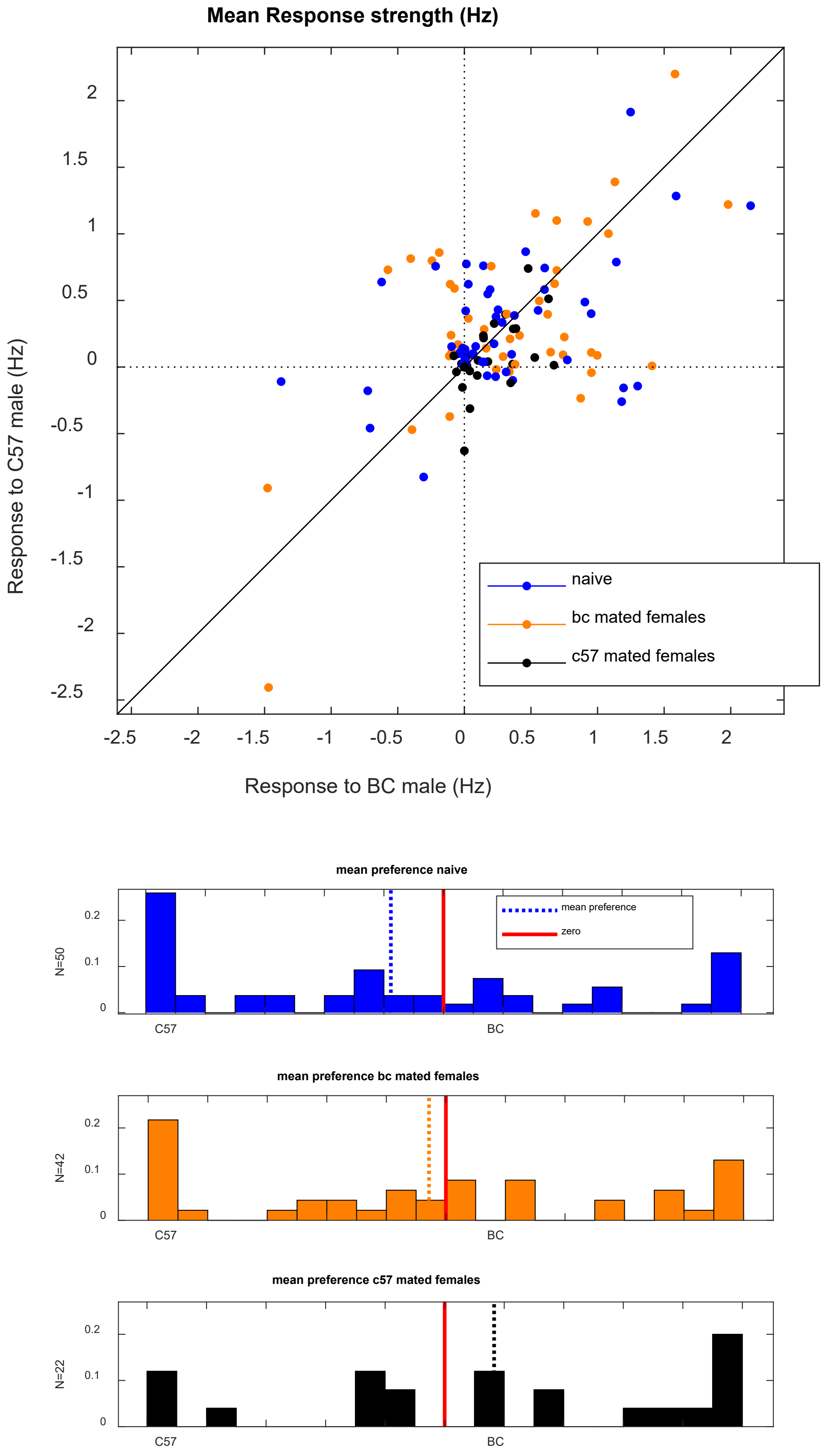


**Figure S6 (part 1). Analysis of response strength to male stimuli from both male strains when response strength is characterized as the average rather than the maximal response. single unit data.** With the exception of this change in quantification, and some formatting changes, the top and bottom panels are analogous to those shown in **Figure 4B** and **Figure 4C**, respectively.

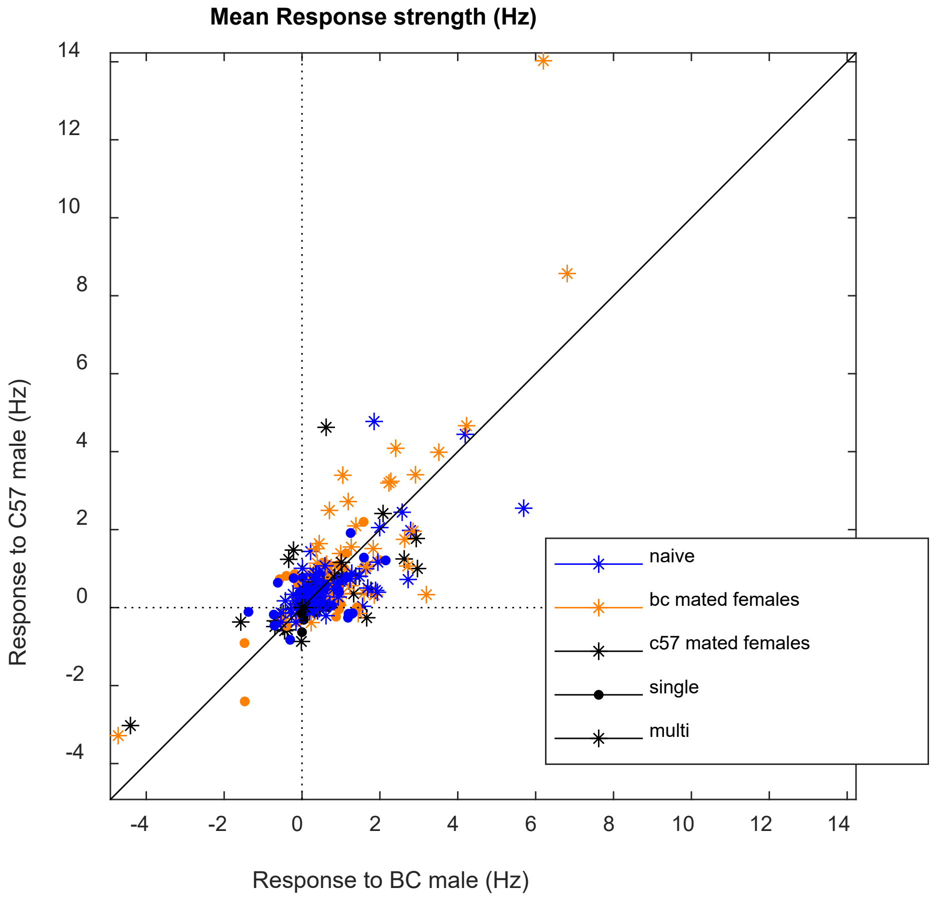


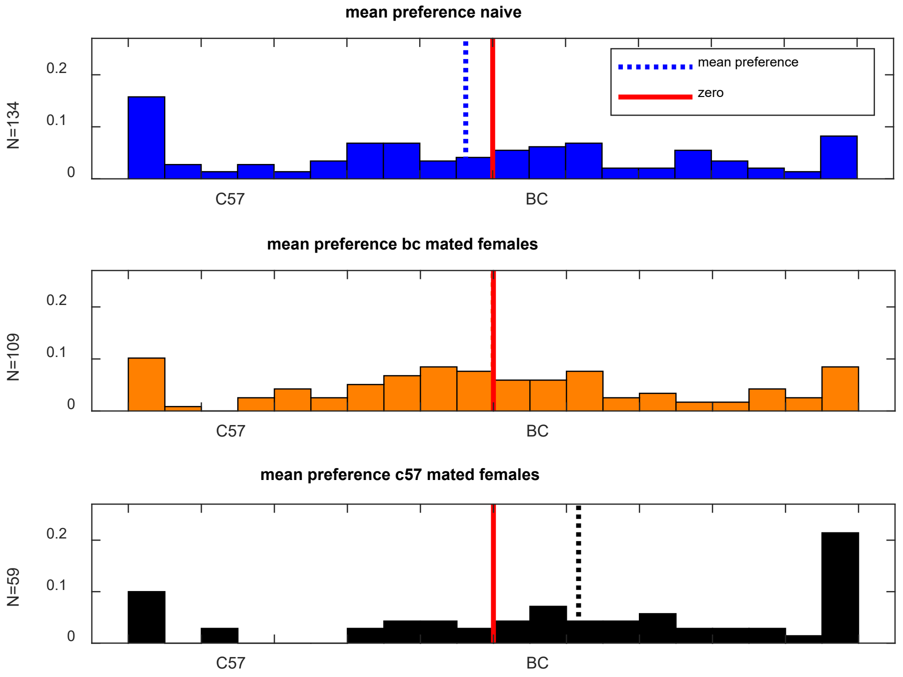


**Figure S6 (part 2). Analysis of response strength to male stimuli from both male strains when response strength is characterized as the average rather than the maximal response. single unit and multi unit data.** With the exception of this change in quantification, and some formatting changes, the top and bottom panels are analogous to those shown in **Figure 4B** and **Figure 4C**, respectively.

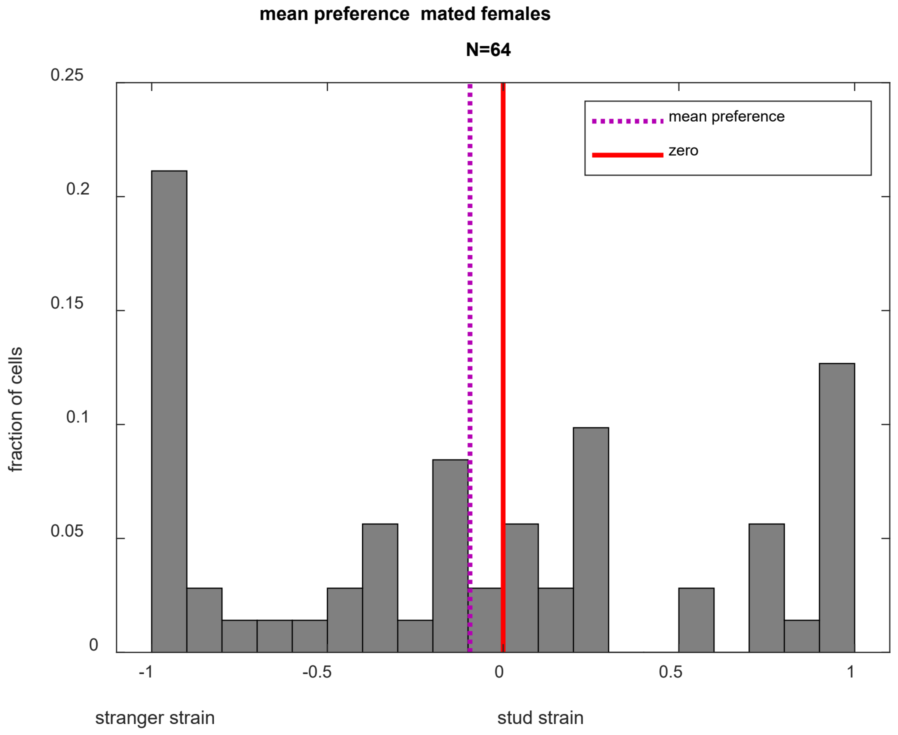


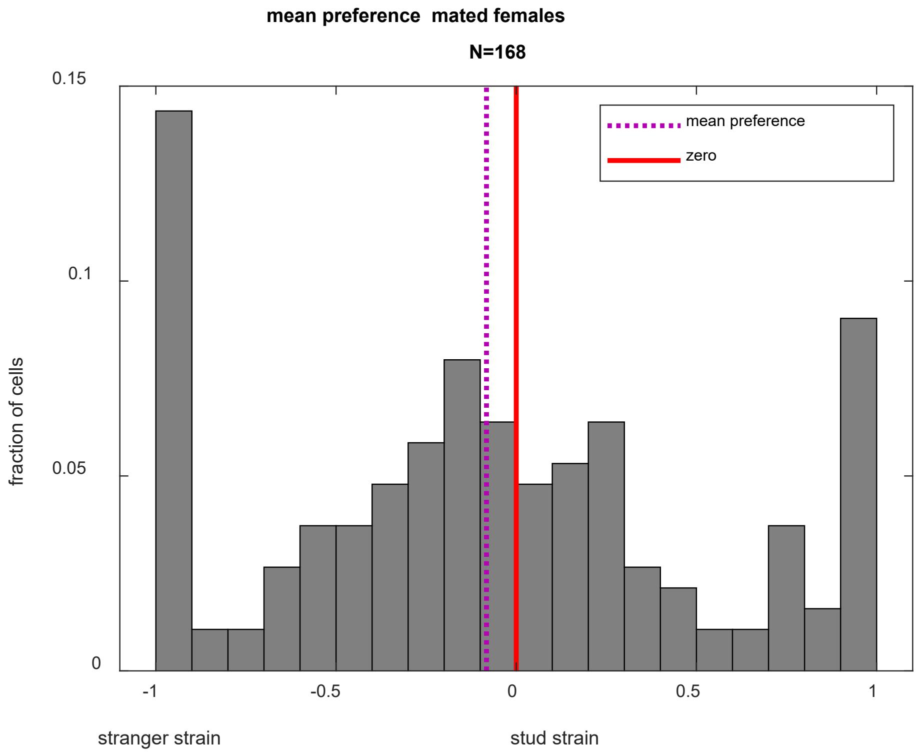


**Figure S6 (part 3). Analysis of response strength to the stud strain stranger strain using the mean rather than maximal response to characterize each response. Top: single unit. Bottom: Single and multi unit.** With the exception of some formatting changes, the top and bottom panels are entirely analogous to those shown in **Figure 4D**. Top: mean index-: -0.0943) , pval (sign rank test): 0.2453. Bottom: mean index: -0.0826), pval (sign rank test): 0.058

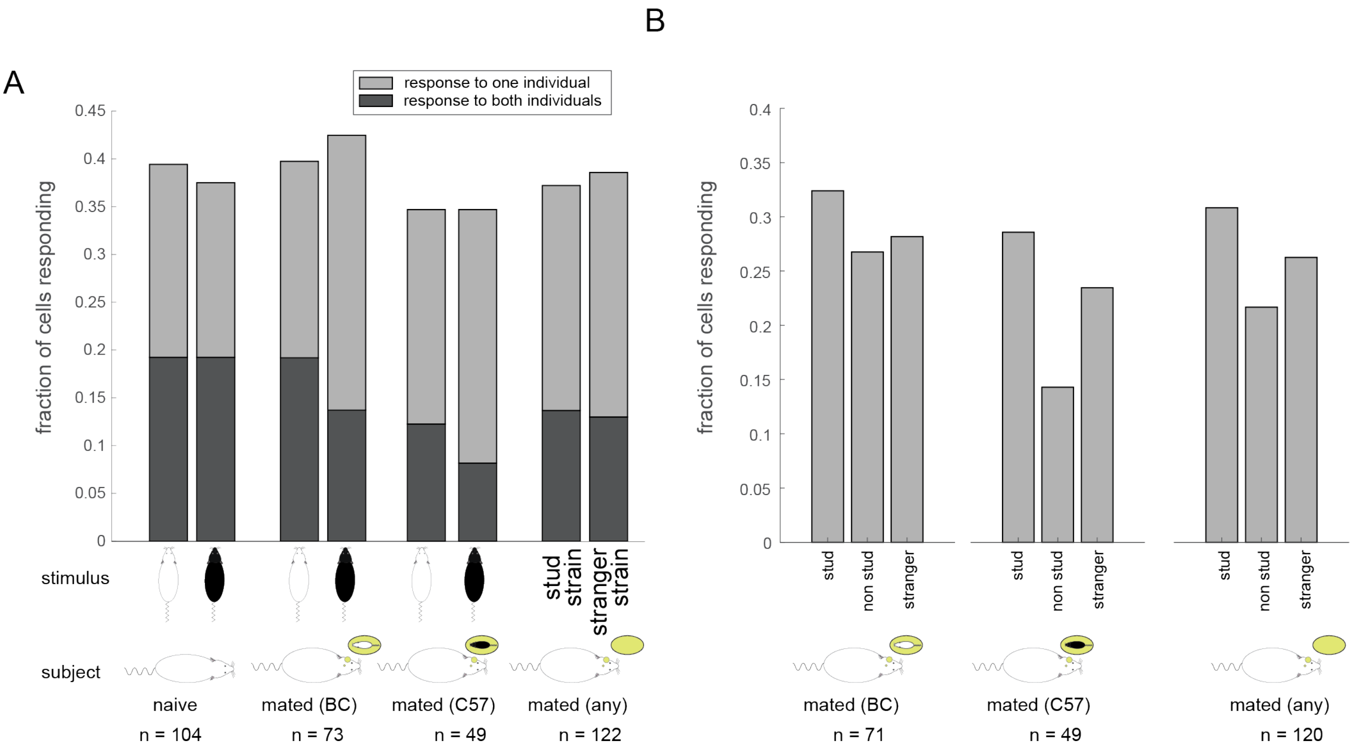


**Figure S7. Distribution of significant responses to male stimuli in naive and mated females. A.** Fraction of single units with significant responses to one or both individuals from the two male strains in the different female groups. **B.** Fraction of single units with responses to the stud individual, the non-stud individual or one of the stranger male individuals. The proportions of neurons that respond to the stud strain or stud male do not change following mating. See text below for details

Analysis of fraction of responding unit (**Figure S7**):

In females mated with BC males, the fraction of single units responding to both BC males is 0.19 while the fraction responding to only one BC male is 0.20, which were identical to the corresponding values in naive females (also 0.19 and 0.20 respectively). In females mated with C57 males, the fraction of neurons responding to both C57 males is 0.08 while the fraction responding to only one C57 male is 0.27, which again was not lower than the corresponding values in naive females (0.19 and 0.18 respectively; **Figure S7A)**. Similar conclusions are reached when combining data from both mated female groups (**Figure S7A**). In this case, the fraction of single units responding to both individuals from the stud male strain is 0.14, similar to the fraction of neurons responding to both individuals from the stranger strain is 0.13. The fraction of neurons responding to one individual from the stud strain is 0.24, also similar to the fraction of neurons responding to one individual from the stranger strain is 0.26 (**Figure S7A)**. None of the groups are statistically distinguishable; comparison of the fraction of responding neurons to one or both individuals between the two male strains is not significantly different ($p>0.05$ using a two-tailed chi-square test of proportions). Taken together, when considering the basic unit as the “strain”, we find no evidence for a reduction in the number of neurons that show a significant *initial* response to one or both individuals from the strain. (continued below ...)

In **Figure S7A**, we do not distinguish among the two individuals from the stud strain. However, it may well be that response silencing will be specific to the stud male rather than to all individuals from that strain. We therefore next asked whether mating selectively silences responses to the stud individual. Here too, we find no evidence for silencing (**Figure S7B**). Specifically, in females that mated with BC males, the fraction of single neurons responding to either the stud or non-stud individual were highly similar (0.32 vs. 0.27). The fraction of neurons responding to individuals from the unfamiliar strain (i.e. C57), averaged across both individuals, is 0.28. In females mated with C57 males, the fraction of neurons responding to the stud individual is 0.29 while the fraction of neurons responding to the non-stud individual from the same strain is 0.14. The fraction of neurons responding to BC individuals (the unfamiliar strain), averaged across the two individuals, is 0.23. Combining the two female groups, the fraction of stud individual responsive neurons is 0.3, while the fraction responding to the same-strain non-stud individual is 0.22. The fraction of neurons responding to the unfamiliar strain (averaged across the two individuals), is 0.26. In neither C57 mated females, BC mated females, or both female groups combined, is there a significant difference in the fraction of stud male responding vs. non-stud male responding neurons ($p>0.05$, chi-square test for proportions). The main conclusions of the analyses remain when multi-units are included (data not shown).

| FB | FC | MB1 | MB2 | MC1 | MC2 | Pred | cast B | cast C |
|---|---|---|---|---|---|---|---|---|
| 0.0002 | 0.1386 | 0.1357 | 0.1887 | 0.5663 | 0.5921 | 0.2556 | 0.0497 | 0.7758 |

**Table S8. Comparison of median response magnitudes between naive and mated females.** P values of two samples non-parametric test (rank-sum) for comparison of each odor response magnitude between naïve and mated females (see Figure 5D).

To obtain the significance threshold of 0.05 after multiple comparisons, we divide by the number of stimuli (9), reaching a threshold of 0.0056. Using this correction (and in fact, also without it), the only significant difference is for the BC female stimulus.

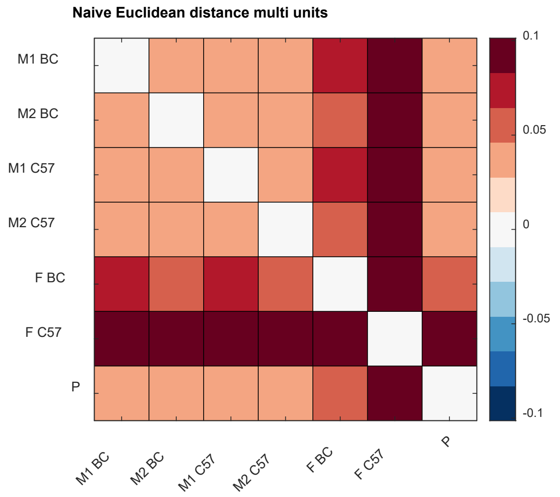


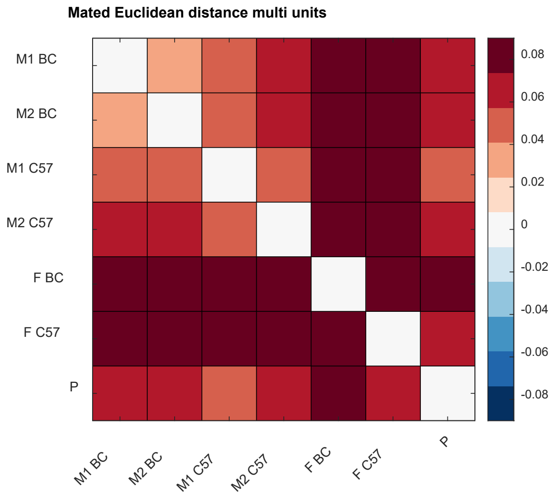


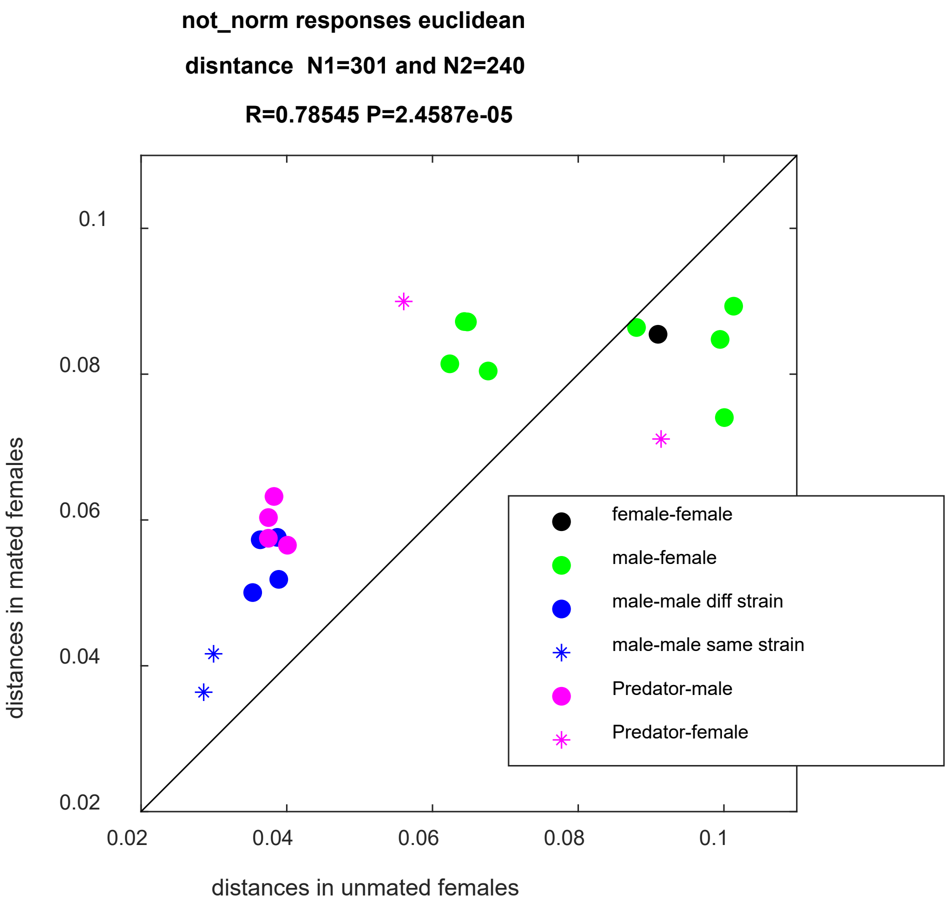


**Figure S9 (part 1). Comparison of pairwise population level distances including multi unit data.** Distance for naive (upper left, n=240 units), mated (upper right, n=301) and their correlation (bottom). Same layout as in Figure 6A-C with some formatting differences. The correlation between the distances and its significance are indicated on title of the bottom plot.

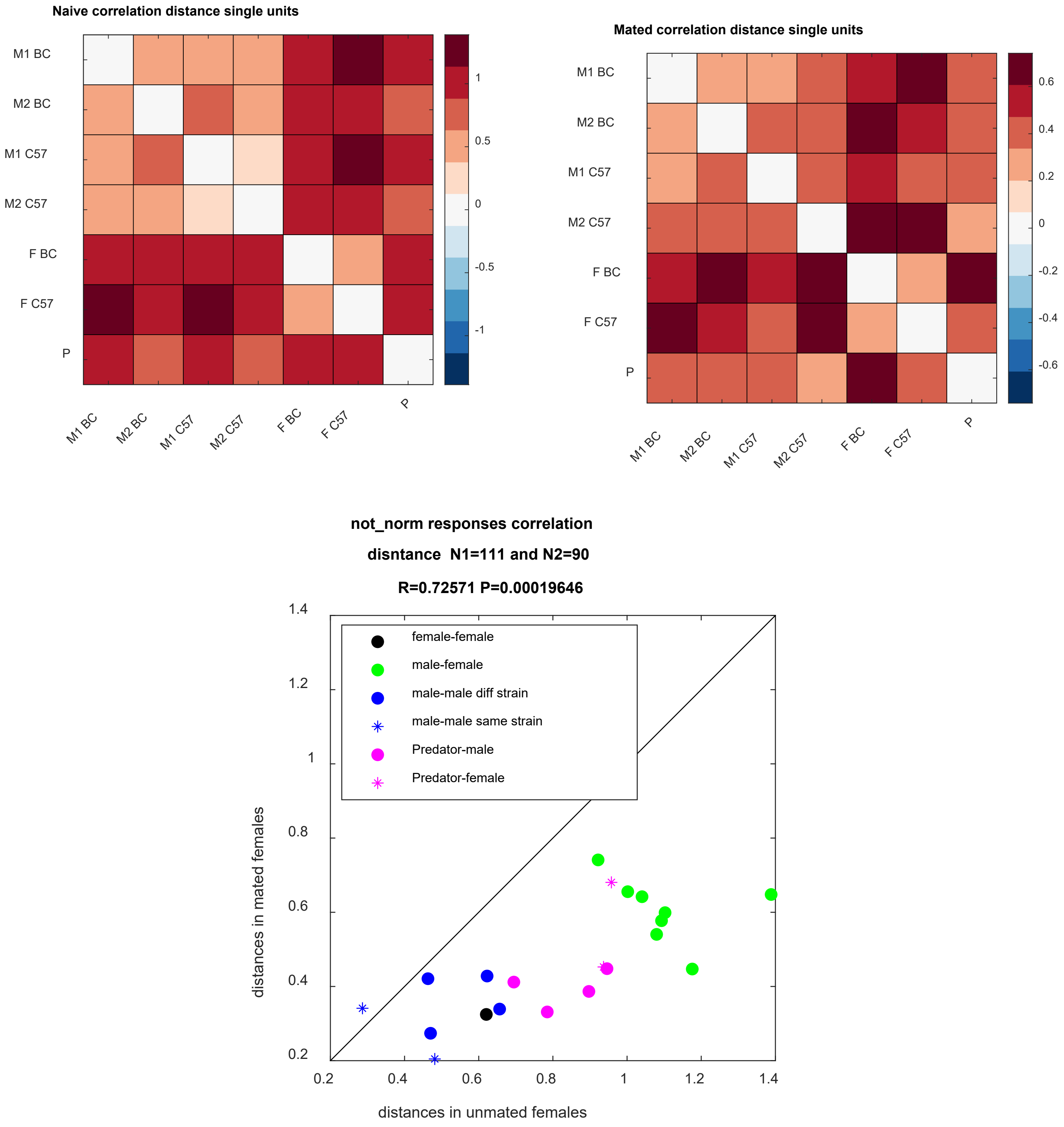


**Figure S9 (part 2). Comparison of pairwise population level distances using the correlation distance measure.** Distance for naive (upper left, n=90 units), mated (upper right, n=111) and their correlation (bottom). Same layout as in Figure 6A-C with some formatting differences. The correlation between the distances and its significance are indicated on title of the bottom plot. The correlation distance measure is defined as one minus the Pearson correlation coefficient.

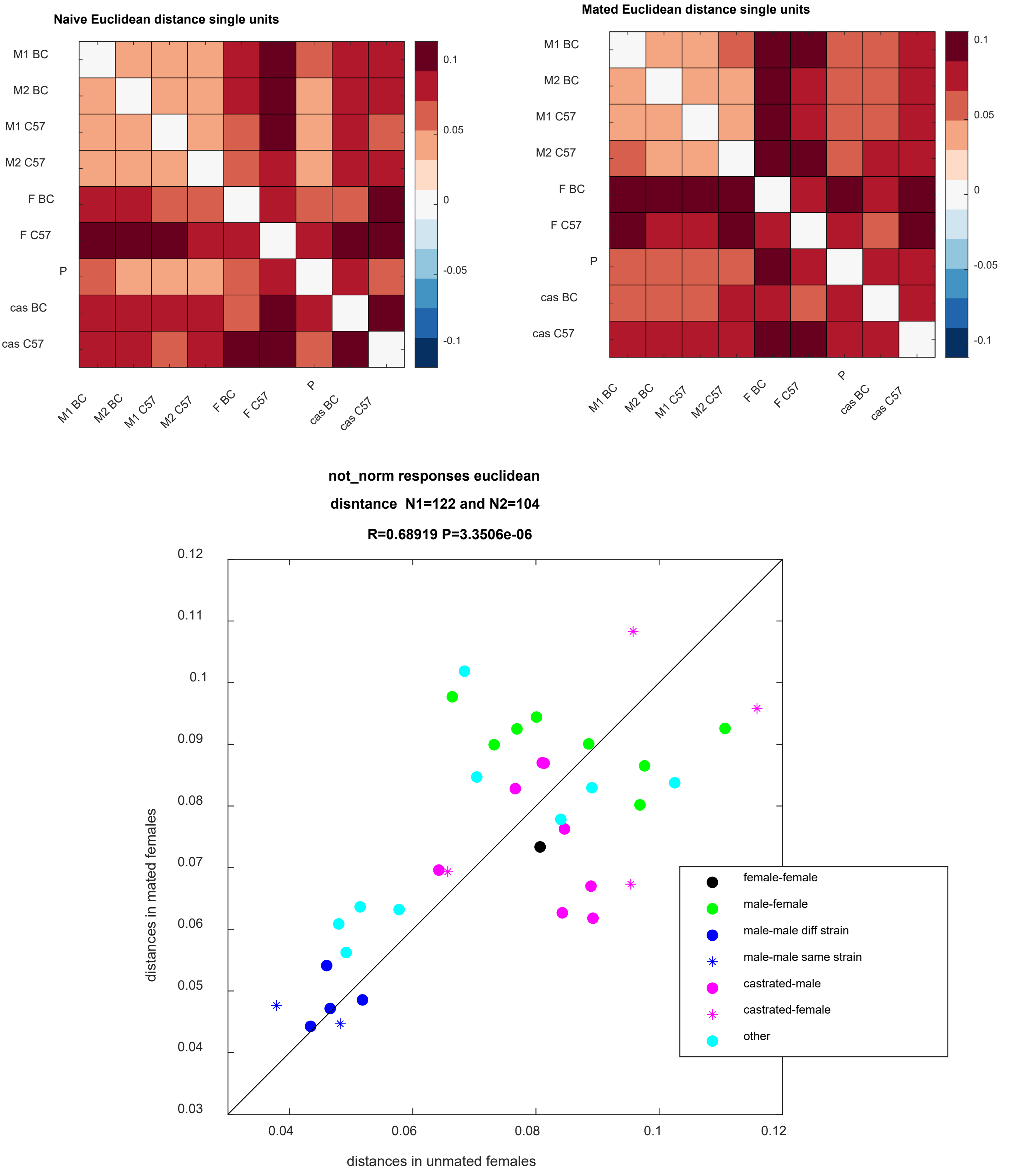


**Figure S9 (part 3). Comparison of pairwise population level distances with inclusion of castrated male stimuli (using single units only and the Euclidean distance measure).** Distance for naive (upper left, n=104 units), mated (upper right, n=122) and their correlation (bottom). Same layout as in Figure 6A-C with some formatting differences. The correlation between the distances and its significance are indicated on title of the bottom plot.
